# A revised perspective on the evolution of troponin I and troponin T in vertebrates

**DOI:** 10.1101/2022.04.22.489046

**Authors:** William Joyce, Daniel M. Ripley, Todd Gillis, Amanda Coward Black, Holly A. Shiels, Federico G. Hoffmann

## Abstract

The troponin (Tn) complex, responsible for the Ca^2+^ activation of striated muscle, is composed of three interacting protein subunits: TnC, TnI, and TnT, encoded by *TNNC*, *TNNI*, and *TNNT* genes. *TNNI* and *TNNT* are sister gene families, and in mammals the three *TNNI* paralogs (*TNNI1*, *TNNI2*, *TNNI3*), which encode proteins with tissue-specific expression, are each in close genomic proximity with one of the three *TNNT* paralogs (*TNNT2*, *TNNT3*, *TNNT1*, respectively). It has been widely presumed that all vertebrates broadly possess genes of these same three classes, although earlier work has overlooked jawless fishes (cyclostomes) and cartilaginous fishes (chimaeras, rays and sharks), which are distantly related to other jawed vertebrates. With a new phylogenetic and synteny analysis of a diverse array of vertebrates including these taxonomic groups, we define five distinct *TNNI* classes (*TNNI1*-5), with *TNNI4* and *TNNI5* being only present in non-mammalian vertebrates and typically found in tandem, and four classes of *TNNT* (*TNNT1-4*). These genes are located in four genomic loci that were generated by the 2R whole-genome duplication events. *TNNI3*, encoding ‘cardiac TnI’ in mammals, was independently lost in cartilaginous and ray-finned fishes. Ray-finned fishes predominantly express *TNNI1* in the heart. *TNNI5* is highly expressed in shark hearts and contains an N-terminal extension similar to that of *TNNI3* found in tetrapod hearts. Given that *TNNI3* and *TNNI5* are distantly related, this supports the hypothesis that the N-terminal extension may be an ancestral feature of vertebrate *TNNI* and not an innovation unique to *TNNI3*, as has been commonly believed.

## Introduction

Contraction of striated muscle is initiated when Ca^2+^ binds to the troponin (Tn) complex, which is located, along with tropomyosin, in association with the actin filament of the sarcomere (Solaro & Rarick 1998; van der Velden & Stienen 2019). Tn-Ca^2+^ binding induces a conformational change that moves tropomyosin and allows the formation of actin-myosin cross-bridges which generate contractile force. The Tn complex is composed of three proteins; the Ca^2+^-binding subunit (TnC), the inhibitory subunit (TnI), and the tropomyosin-binding subunit (TnT) (van der Velden & Stienen 2019; Solaro & Rarick 1998). TnC is a calmodulin-like protein that is part of the helix-loop-helix group of Ca^2+^ binding proteins (Strynadka & James 1989), whilst TnI and TnT, which are closely related to one another (Chong & Jin 2009; Rasmussen & Jin 2021; Wei & Jin 2016), indirectly affect Ca^2+^ affinity of Tn through protein-protein interactions within the complex (Lombardi et al. 2008; Evans & Levine 1980; Hwang et al. 2014). This mode of contraction activation is evolutionarily ancient and can be traced back to the earliest bilaterian animals ∼700 million years ago (Barnes et al. 2016; Rasmussen et al. 2022; Cao et al. 2019; Yaguchi et al. 2017)

In vertebrates, each striated muscle type (*i.e.* cardiac muscle, slow and fast twitch skeletal muscle) express a specific complement of TnC, TnI, and TnT genes (*TNNC*, *TNNI* and *TNNT*, respectively). It is widely accepted that in mammals and most other vertebrates, two groups of genes encoding TnC are found, which are characterised by expression in fast skeletal muscle (fsTnC; *TNNC2*) or both cardiac and slow skeletal muscle (cTnC; *TNNC1*) (Gillis et al. 2007). In some ray-finned fishes, including teleosts and gar, there are two *TNNC1* genes due to a lineage-specific gene duplication (Genge et al. 2016). The evolutionary histories of TnI and TnT have received great attention (Wei & Jin 2016; Rasmussen et al. 2022; Sheng & Jin 2016; MacLean et al. 1997; Palpant et al. 2010; Gross & Lehman 2016; Chong & Jin 2009; Hastings 1997; Shaffer & Gillis 2010; Genge et al. 2016; Barnes et al. 2016) and it is generally believed that, like mammals, most vertebrates possess three genes encoding TnI; slow skeletal (ssTnI; *TNNI1*), fast skeletal (fsTnI; *TNNI2*) and cardiac (cTnI; *TNNI3*), and three genes for TnT; slow skeletal (ssTnT: *TNNT1*), fast skeletal (fsTnT; *TNNT3*) and cardiac (cTnT: *TNNT2*). Because we explore a range of previously uncharacterised *TNNI* and *TNNT* genes and proteins outside of the three defined in mammals, and because non-mammalian vertebrates are known to express a variety of *TNNI*s in a given muscle type (Alderman et al. 2012), we herein eschew the protein names that derive from muscle specific expression and instead adopt protein names based on the corresponding numbered gene *i.e.* TnI1-3 corresponding to genes *TNNI1-3* (instead of ssTnI, fsTnI and cTnI respectively).

Mammalian *TNNI* genes are located in close proximity to *TNNT* paralogs in human and mouse genomes: *TNNI2* with *TNNT3*, *TNNI3* with *TNNT1*, and *TNNI1* with *TNNT2* (Chong & Jin 2009) and there is also some limited evidence for this in fish (Genge et al. 2016). This is intriguing because whole-genome duplications (WGDs) have played a key role in expanding the gene repertoireof early vertebrates (Hoffmann et al. 2021, 2012; Zavala et al. 2017), and because the teleost whole-genome duplication has been linked to the functional diversification of the zebrafish *TNNI* paralogs (Genge et al. 2016). As such, the diversity in *TNNI* and *TNNT* families appears to have arisen through a tandem duplication followed by successive rounds of whole-genome duplication (Chong & Jin 2009; Rasmussen & Jin 2021).

Particular interest has been paid to the evolution of *TNNI3*, which in adult mammals is solely expressed in the heart and is distinguished from other vertebrate *TNNI* paralogs by a ‘unique’ N-terminal extension peptide (Sheng & Jin 2016; Shaffer & Gillis 2010; Rasmussen et al. 2022). This N-terminal extension is an important regulatory structure (Sheng & Jin 2016) containing two protein kinase A (PKA) target serine residues that, when phosphorylated via ß-adrenergic stimulation, decrease myofilament Ca^2+^ sensitivity (Fentzke et al. 1999; Layland et al. 2005; Robertson et al. 1982, 198; Solaro et al. 1976, 197), thereby increasing the rate of relaxation during diastole (Kentish et al. 2001; Zhang et al. 1995). In teleost fishes, such as zebrafish (Fu et al. 2009) and rainbow trout (Alderman et al. 2012; Gillis & Klaiman 2011; Kirkpatrick et al. 2011), cardiac-expressed TnI lacks the N-terminal extension that characterises mammalian TnI3, although a long N-terminal extension is present in amphibian (Drysdale et al. 1994; Warkman & Atkinson 2004) and lungfish (Rasmussen et al. 2022) TnI3. Currently, the most widely believed consensus, at least in the vertebrate TnI field, is that *TNNI1* and *TNNI3* are more closely related to each other than to *TNNI2*, and that all three evolved from a single gene in the ancestor of vertebrates (Shaffer & Gillis 2010; Sheng & Jin 2016). The N-terminal extension has been commonly interpreted as an evolutionary novelty that emerged in TnI3 from a TnI1-like ancestral form in the ancestor of lobe-finned fishes (Palpant et al. 2010; Shaffer & Gillis 2010; Sheng & Jin 2016; Rasmussen et al. 2022).

However, given that some protostome invertebrate (Cao et al. 2019; Barnes et al. 2016) and tunicate (MacLean et al. 1997) *TNNI* genes encode for alternatively-spliced isoforms with and without an N-terminal extension, an alternative interpretation is that skeletal muscle paralogs secondarily lost an ancestral extension that has been differentially retained by*TNNI3* (Hastings 1997; MacLean et al. 1997; Barnes et al. 2016).

Several fundamental questions remain open regarding the evolution of the troponin I and T genes in vertebrates. Both teleost fish and mammals have *TNNI* paralogs that encode for cardiac-expressed TnIs, however, it is not clear whether these subunits are encoded by orthologous genes (Genge et al. 2016; Rasmussen et al. 2022). Further, the duplicative history of these genes in the early stages of vertebrate evolution could not be properly resolved because of the limited availability of cartilaginous fish and jawless fish sequences. Cartilaginous fish sequences are particularly valuable as this lineage was the first to diverge from other gnathostomes (jawed vertebrates), meaning that orthologous genes identified in both cartilaginous fishes and other gnathostome clades can be traced back to the last common ancestor of all jawed vertebrates. The current consensus is that the three tetrapod *TNNI*s are monophyletic, and that tetrapod and ray-finned fishes *TNNT2*s fall in a monophyletic group, implying these genes are orthologs. The evidence is inconclusive regarding *TNNI1* and *TNNI3* (Genge et al. 2016, Sheng & Jin 2016). Support to resolve relationships among the different vertebrate *TNNI* paralogs is also limited (Genge et al. 2016, Sheng & Jin 2016), which is critical to determine whether the N-terminal extension of cTnI is ancestral or derived, and to provide a robust evolutionary context to interpret the observed functional differences among the different TnI subunits, including the capacity to regulate contractile function via ß-adrenergic stimulation.

In the current study, we take advantage of improved assemblies of sea lamprey and cartilaginous fish genomes to answer long-standing questions on the duplicative history of the *TNNI* and *TNNT* gene families of vertebrates. We combine phylogenetic and synteny analyses from a representative set of vertebrates to reconstruct the early stages of evolution of these two closely related gene families in the group. Our reconstruction indicates that the last common ancestor of gnathostomes possessed five *TNNI* and four *TNNT* genes in its genome arranged in four different loci which derive from the 2R of WGD early in vertebrate evolution. Comparisons with lamprey and hagfish suggest that the tandem arrangement of *TNNI* and *TNNT* was present in the last common ancestor of vertebrates. We augment our analyses by assessing *TNNI* gene and protein expression, as well as PKA-mediated phosphorylation of cardiac-expressed TnI, in a diverse cohort of gnathostome vertebrates. In the context of our phylogenetic findings, our results indicate that the presence of an N-terminal extension in the TnI3 subunit of tetrapods represents the retention of an ancestral feature rather than an evolutionary innovation of tetrapods or sarcopterygian fish. Our findings also indicate that the genes encoding for the cardiac-expressed TnI subunits of teleost fish (*TNNI1*) and tetrapods (*TNNI3*) are not orthologs. Instead, these subunits are encoded by paralogous genes that were lost (*TNNI3* in ray-finned fish lineage) or exhibit divergent expression patterns (*TNNI1* being restricted to slow skeletal muscle and embryonic cardiac muscle in tetrapods).

## Results

### Data description and approach

We combined bioinformatic searches of the NCBI and Ensembl databases to collect the full *TNNI* and *TNNT* repertoires in a representative set of vertebrate genomes including two invertebrate chordates as reference. Because our aim was on the early stages of vertebrate evolution and on resolving relationships between teleost and tetrapod paralogs, our sampling included a focussed number of amniotes and teleosts. Our set consisted of 19 different species that included two cyclostomes (Sea lamprey, *Petromyzon marinus* and Inshore hagfish, *Eptatretus burgeri*); representatives of three different orders of cartilaginous fishes (class Chondrichthyes), elephant fish, also known as elephant shark (*Callorhinchus milii*, order Chimaeriformes), thorny skate (*Amblyraja radiata,* order Rajiformes), and small-spotted catshark (*Scyliorhinus canicula*, order Carcharhiniformes); three non-teleost ray finned fishes, reedfish (*Erpetoichthys calabaricus,* order Polypteriformes), sterlet sturgeon (*Acipenser ruthenus*, order Acipenseriformes) and spotted gar (*Lepisosteus oculatus*, order Lepisosteiformes); two teleosts, Asian bonytongue (*Scleropages formosus*, order Osteoglossiformes), and zebrafish (*Danio rerio*, order Cypriniformes); African coelacanth (*Latimeria chalumnae,* order Coelacanthiformes); West African lungfish (*Protopterus annectens,* order Dipnoi); an amphibian, tropical clawed frog (*Silurana tropicalis,* order Anura); a non-avian reptile, anole lizard (*Anolis carolinensis*, order Squamata); a bird, chicken (*Gallus gallus*, order Galliformes); a monotreme, Australian echidna (*Tachyglossus aculeatus*, order Monotremata); and a eutherian mammal, human (*Homo sapiens*, order Primates) (databases available in Supplementary Material online). As outgroup references, we included the full repertoire of *TNNI* and *TNNT* genes from two invertebrate chordates: the sea squirt (*Ciona intestinalis*, a tunicate), and the Florida lancelet (*Branchiostoma floridae*, a cephalochordate). Because we studied a broad range of taxonomic groups with different gene nomenclature practises (*e.g.* teleost “*tnni*”), to avoid confusion we standardised all gene names to the human convention *e.g.* “*TNNI*”.

Our bioinformatic searches combined information from the Ensembl comparative genomics assignments of orthology (Zerbino et al. 2018) with the results of BLAST searches (NCBI Resource Coordinators 2016) against the corresponding genomes. BLAST searches used the blastp and tblastn programs and were seeded with known *TNNI* and *TNNT* protein sequences. We validated our *TNNI* and *TNNT* candidates using reverse BLAST against the NCBI Reference Protein database of vertebrates, refseq_protein. Candidate records that did not include either a *TNNI* or *TNNT* as their top hit were discarded. We inferred that our sequences had captured the full range of *TNNI* and *TNNT* diversity present in each of the genomes surveyed because searches seeded with *TNNI* identified *TNNT*-like sequences, and searches seeded with *TNNT* sequences identified *TNNI*-like sequences.

### Variation in TNNI and TNNT gene complements

After curating the results of our searches, our data sets included a total of 81 *TNNI* and 72 *TNNT* sequences. As expected, vertebrates exhibit a wider range of variation in both gene families relative to the invertebrate chordates included as outgroups. In the case of *TNNI*, the number of genes in invertebrate chordates ranged from one in sea squirt (a tunicate) to two in the Florida lancelet (amphioxus), whereas in vertebrates the number ranged from two in reedfish, the least of any vertebrate we surveyed, to a maximum of 14 in zebrafish, which have undergone an additional WGD, and include two series of tandem duplications. In general, our results agree with previous assessments of copy number variation for this gene family in invertebrate chordates and vertebrates (Shih et al. 2015; MacLean et al. 1997). In the case of the *TNNT* genes, the number of genes in invertebrate chordates matched the number of *TNNI* genes, one in sea squirt and two in the Florida lancelet, and in the case of vertebrates, the number ranged from three in gar, coelacanth, and tetrapods to the eight different copies identified in zebrafish.

### Phylogenies identify additional gnathostome *TNNI* and *TNNT* paralogs

Our phylogenetic analyses place vertebrate *TNNI*s in a monophyletic clade and arrange gnathostome *TNNI*s into five strongly supported monophyletic groups (fig. 1). Three of these groups can be defined by the presence of the mammalian *TNNI*1, *TNNI*2, and *TNNI*3 paralogs. The *TNNI1* and TNNI2 groups include ray-finned, lobe-finned, and cartilaginous fish genes, whereas we only found *TNNI3* copies in lobe-finned fishes. The fourth group, *TNNI4*, contains previously annotated (*i.e.* NCBI and ZFIN) *tnni4* zebrafish genes, although these have not previously been formally described in the literature. For the remaining group we coin the name *TNNI5*. *TNNI4* and *TNNI5*, have restricted phyletic distributions, and we failed to find copies of these genes in any of the amniote genomes we surveyed. *TNNI4* is present in the genomes of cartilaginous fishes, ray-finned fishes, lungfish, and amphibians, and includes the previously identified *tnni1.2* gene of the tropical clawed frog (NCBI Gene ID: 394556; Xenbase: XB-GENE-485710, Table 1). In turn, *TNNI5* is restricted to cartilaginous fishes, ray-finned fishes, and coelacanth, and includes the previously named zebrafish genes *tnni1c* (NCBI Gene ID: 751665, ZFIN:ZDB-GENE-060825-192, Table 1) and *tnni1d* (NCBI Gene ID: 436902, ZFIN:ZDB-GENE-040718-374, Table 1). Our analyses identify additional duplications, found in lineages that have undergone additional WGDs, such as teleosts or sterlet, some of which correspond to the already reported tandem expansions of *TNNI2* in zebrafish.

**Figure 1.**
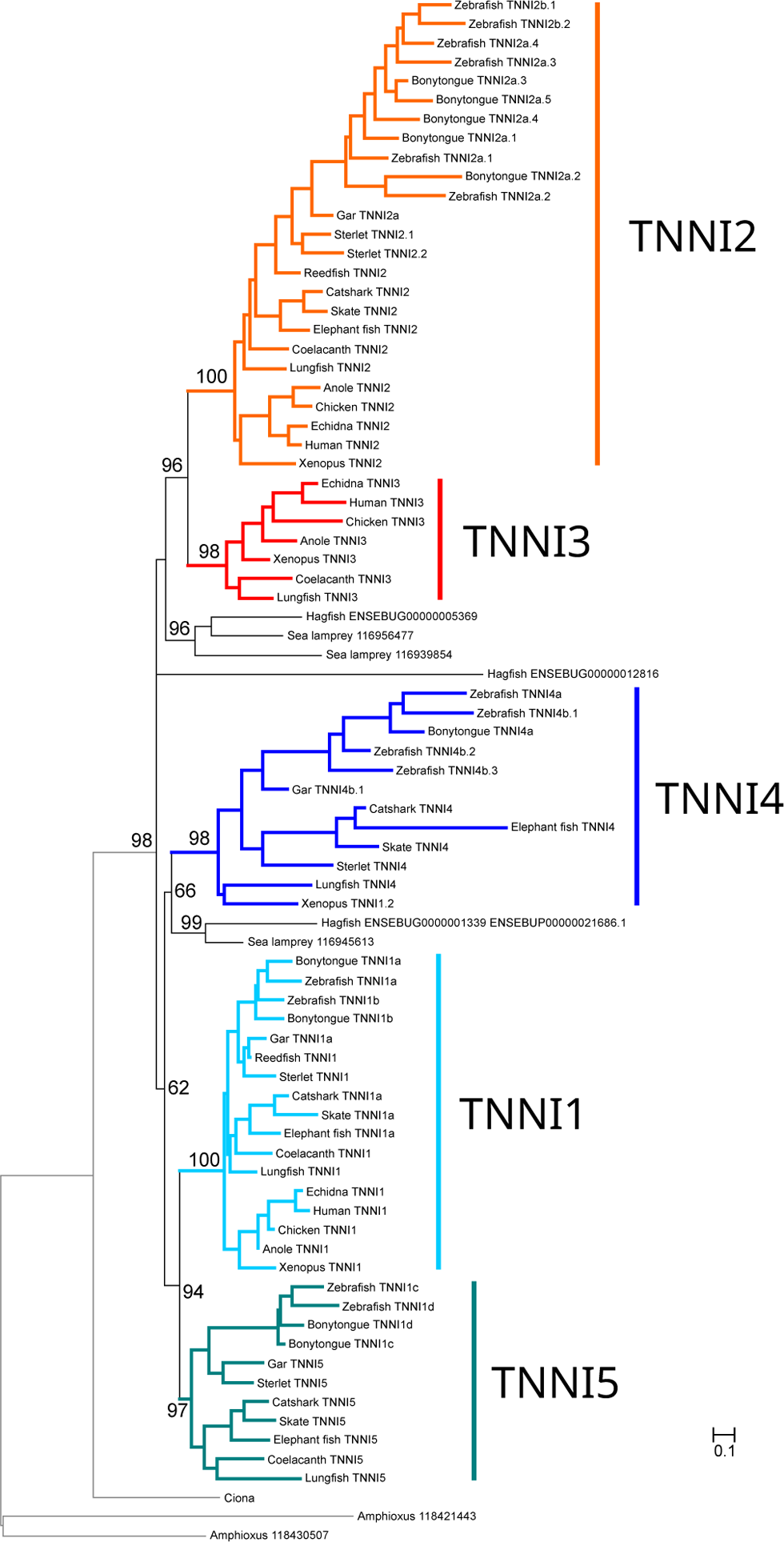
Maximum likelihood phylogenetic tree showing evolutionary relationships between vertebrate troponin I (*TNNI*) sequences. The five distinct *TNNI* groups that we infer were present in the common ancestor of gnathostome vertebrates are highlighted. The tree was rooted with the amphioxus and Ciona sequences. Ultrafast bootstrap support is shown above relevant nodes.

**Table 1.**
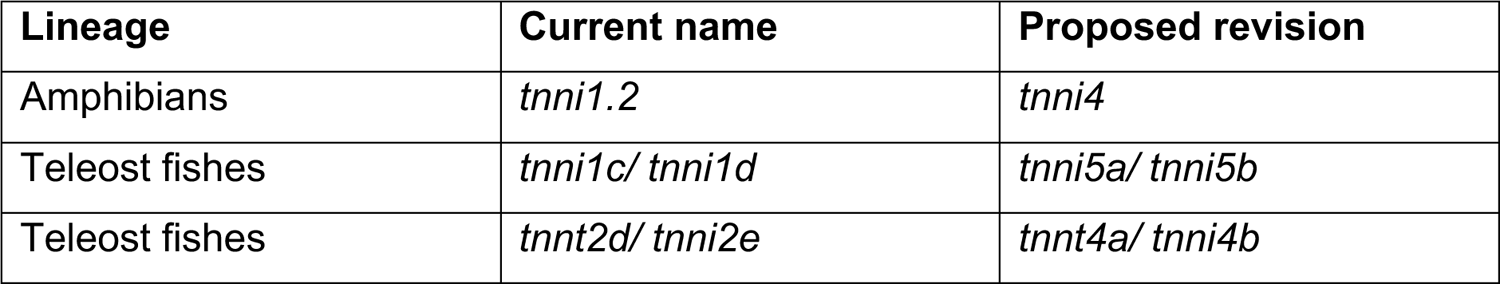
Proposed changes in nomenclature for previously mis-annotated genes. Revised proposed names are intended to reflect the evolutionary history and phylogenetic affiliations within newly defined *TNNI* and *TNNT* groups.

The 5 gnathostome *TNNI* genes are divided into two super-groups, *TNNI3* is placed as sister to *TNNI2* in the first group, and with *TNNI1, 4*, and *5* are placed in the second one, where *TNNI1* is placed as sister to *TNNI5*, and *TNNI4* groups with the *TNNI1*+TNNI5 clade (fig. 1). Lamprey and hagfish include 3 *TNNI* paralogs in their genomes that broadly clustered with the gnathostome *TNNI2/3* or *TNNI1/4/5* groups. The lamprey *116956477* and *116939854* genes are placed in a group with the hagfish *ENSEBUG00000005369* gene in a clade that is placed within gnathostome *TNNI2/3*, and the lamprey *116945613* gene is sister to the hagfish *ENSEBUG00000013390* in a clade within the *TNNI1/4/5 clade of gnathostomes*. The third hagfish *TNNI*, *ENSEBUG00000012816*, appears to be either incomplete in the current genome annotation or highly divergent and has unclear phylogenetic affinities.

Like *TNNI*s, vertebrate *TNNT*s were monophyletic relative to invertebrate chordates, and the gnathostome sequences were arranged into four monophyletic groups with moderate to strong support (fig. 2). The *TNNT1*, *TNNT2*, and *TNNT3* groups can be defined by the presence of human paralogs and are found in the vast majority of species surveyed. The fourth group, which we label as *TNNT4* is restricted to cartilaginous fishes, ray-finned fishes and lungfish. As with *TNNIs*, our analyses identify additional duplications which are mostly restricted to lineages that have undergone additional WGDs, such as teleosts or sterlet. The only exception to this is the presence of duplicate *TNNT2*s in reedfish, sterlet and zebrafish. Within the *TNNT2* paralog, tetrapod and cartilaginous fish sequences fell in monophyletic clades, but ray-finned fish sequences were grouped in two separate groups, the first one including copies from all ray-finned fish species in our study, and the second one including copies from reedfish, sterlet, and zebrafish. This arrangement is suggestive of an old duplication in the last common ancestor of ray-finned fishes that has been differentially retained in some descendants. The *TNNT* genes of gnathostomes are arranged in two groups, with *TNNT2* and *TNNT4* in the first, and *TNNT1* and *TNNT3* in the second.

**Figure 2.**
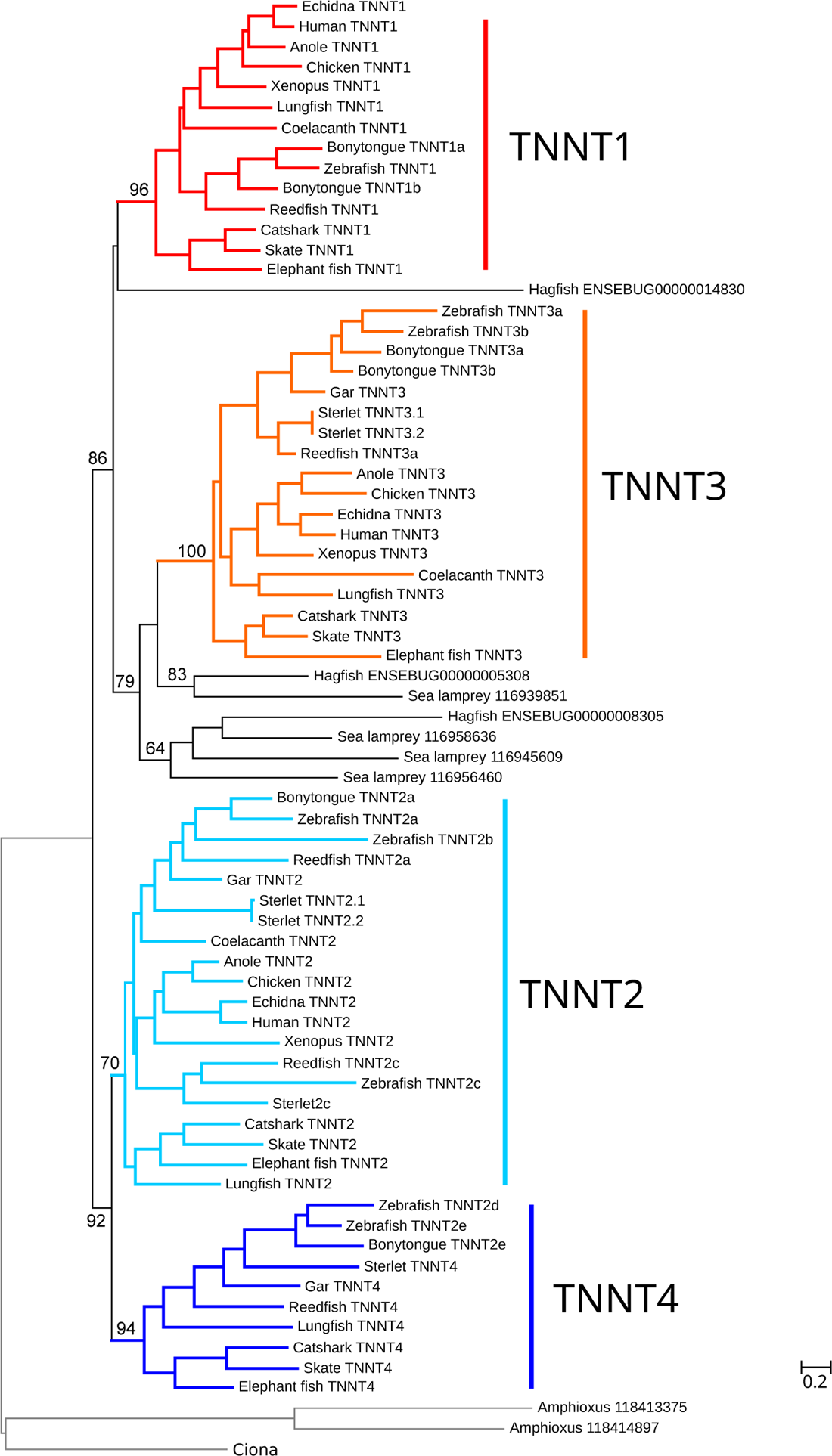
Maximum likelihood phylogenetic tree showing evolutionary relationships between vertebrate troponin T (*TNNT*) sequences. The four distinct *TNNT* groups that we infer were present in the common ancestor of gnathostome vertebrates are highlighted. The tree was rooted with the amphioxus and Ciona sequences. Ultrafast bootstrap support is shown above relevant nodes.

Cyclostomes also include multiple *TNNT* copies in their genomes: four in the case of lamprey and three in the case of hagfish. These genes are arranged into three separate groups, all of which fall within the *TNNT1/3* clade of gnathostomes. The hagfish ENSEBUG00000014830 gene is placed as sister to *TNNT1* with low support. Then a clade with the hagfish ENSEBUG00000005308 and lamprey 116939851 genes is placed as sister to the *TNNT3* group of gnathostomes, and a second cyclostome *TNNT* clade which includes the hagfish ENSEBUG00000008305 gene plus the three remaining lamprey genes, 116945609, 116956460, and 116958636, is placed with gnathostome *TNNT3* as well. Support for the nodes resolving affinities for these groups is low.

### *TNNI*s and *TNNT*s are found in clusters of conserved synteny

In the case of gnathostomes, the results of our synteny analyses (fig. 3) are consistent with our phylogenetic analyses and provide additional insights regarding the duplicative history of the gene families and the absence of some paralogs in some gnathostome genomes. Microsynteny is very conserved in the cases of the *TNNI1-TNNT2* and *TNNI2-TNNT3 clusters of* gnathostomes (fig. 3). *TNNI1* and *TNNT2* are found in tandem, with *LAD1* between them and *PHLDA3* and *PKP* flanking the cluster. *TNNI2* is found in tandem with *TNNT3*, with copies of *LSP1* and *PRR33* between them, and copies of *SYT8* upstream of the cluster. *TNNI3* is flanked by copies of *DNAAF3* and *TNNT1* in most tetrapods. Interestingly, *DNAAF3* and *TNNT1* are adjacent to each other in the elephant fish and reedfish genomes, whereas we could not find copies of any of these three genes in the current release of the spotted gar genome (assembly name: LepOcu1; accession GCF_000242695.1). *TNNI4* and *TNNI5* are found in tandem in cartilaginous fishes and sterlet but are located on separate loci in gar and zebrafish. There are copies of CALD1 and BPGM between *TNNT4* and the *TNNI4-5* cluster. In gar and zebrafish, *TNNI5* is on a different chromosome than the *TNNI4-TNNT4* cluster, flanked by copies of *B4GALNT3B* and *C2CD5*. Orthologs of these two genes are found on separate chromosomes in humans. Further, the arrangement of *TNNI* and *TNNT* genes in the phylogenies is consistent with their position in the genome. The *TNNI2* and *TNNT3* genes, which are found on the same locus, on human chromosome 11, are grouped with *TNNI3* and *TNNT1*, which are found on the same locus, on human chromosome 19. In turn, *TNNI1* and *TNNT3*, which are found on the same locus, on human chromosome 1, are grouped with *TNNI4+TNNI5* and *TNNT4* respectively, which are absent in humans, but are found in the same locus in cartilaginous fish, and sterlet.

**Figure 3.**
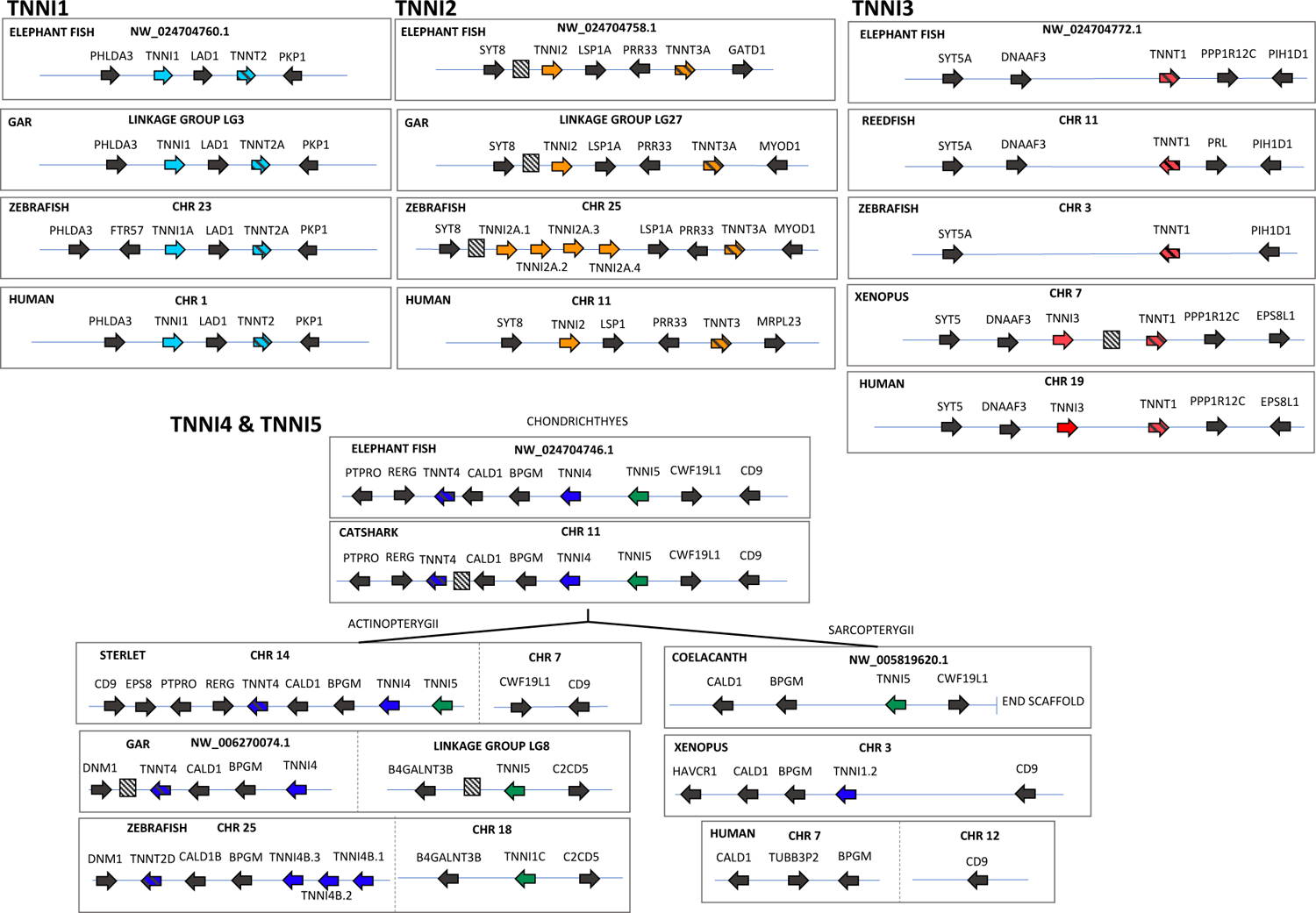
Conserved synteny diagrams of genomic regions harbouring *TNNI*/*TNNT* genes in gnathostome vertebrates. Hatched boxes indicate uncharacterised predicted protein coding genes. Non-coding and microRNAs are excluded.

As in gnathostomes, we find pairs of *TNNT* and *TNNI* genes in close proximity in both cyclostome genomes. However, synteny comparisons are not as informative because of the reduced contiguity of the hagfish genome relative to the lamprey assembly and because there are discrepancies between the phylogenetic and synteny analyses in this group. The lamprey genome includes three *TNNI*-*TNNT* pairs, on chromosomes 7, 24, and 65, plus a single *TNNT* gene on chromosome 80, whereas the hagfish genome contains one pair on contig FYBX02009389. There are copies of *GATD1* and *CALD1* between the *TNNI*-*TNNT* pair on lamprey chromosome 24 (*116945613* and *116945609*) and the hagfish pair on contig FYBX02009389 (*ENSEBUG00000013390* and *ENSEBUG00000005308*). However, the flanking genes are different, and the phylogenies are not congruent with the synteny. The corresponding *TNNI* genes are sister but not the *TNNT* genes. In the amphioxus and the tunicate, *TNNI* and *TNNT* were not found in close genomic proximity, although curiously in amphioxus we found *TNNI* in a cluster with three *TNNC* genes fig. 4).

**Figure 4.**
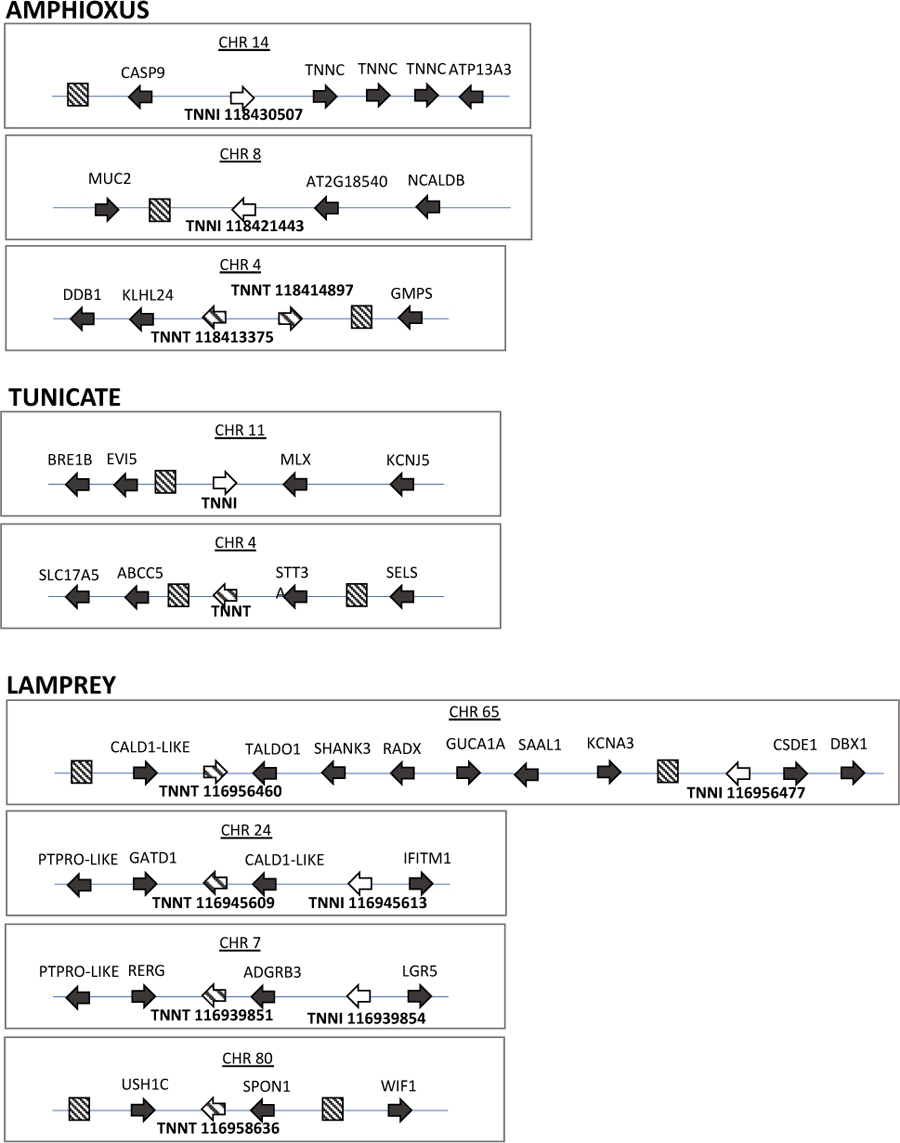
Conserved synteny diagrams of genomic regions harbouring *TNNI*/*TNNT* genes in invertebrate chordates and a cyclostome.

There are similarities in genomic context between the lamprey and the gnathostome *TNNI*-*TNNT* pairs but results of synteny comparisons and phylogenetic analyses are not easy to reconcile. For example, the *TNNT4* genes of cartilaginous fishes and sterlet are next to *PTPRO* and *RERG* copies, as is the *116939851 TNNT* gene of the lamprey, but these genes are not placed together in the phylogeny, and the associated TNNI genes are not placed close together either. More generally, there are similarities in the genomic context shared by many of the *TNNI*-*TNNT* genomic loci. There are paralogs of *CALD1* or *LSP1*, next to two of the *TNNI*-*TNNT* clusters or gnathostomes and two of the lamprey clusters (fig. 4), there are *PTPRO* paralogs close to the *TNNI1*-*TNNT2* and the *TNNI4*-5-*TNNT4* clusters of gnathostomes and the lamprey *TNNI*-*TNNT* clusters on chromosomes 7 and 24, and there are *SYT* paralogs close to the each of the tree *TNNI*-*TNNT* pairs defined by the presence of mammalian *TNNI*s.

### The evolution of the N-terminal extension in TnI

We recognised that the *TNNI5* sequence found in cartilaginous fishes, non-teleost ray-finned fishes, and sarcopterygian fishes included an N-terminal sequence bearing a striking similarity to *TNNI3* previously described in tetrapods (Drysdale et al. 1994; Warkman & Atkinson 2004) and lungfish (Rasmussen et al. 2022).

Although teleost fishes possessed genes of the *TNNI5* family (previously named *tnni1*c and *tnni1d*), they did not contain the N-terminal extension, indicating it was lost in this protein lineage in teleosts. To provide a formal and unbiased comparison between *TNNI* paralogs, we used ancestral sequence reconstructions to predict ancestral protein sequences for *TNNI1-5*. The alignment highlights the strong resemblance between *TNNI3* and *TNNI5* N-terminals, particularly with regard to glutamic acid and proline-rich stretches (fig. 5A). Curiously, the *TNNI3* and *TNNI5* N-terminal extensions also showed similarities to *TNNT*, as represented by a reconstruction of the common ancestor of all gnathostome vertebrate *TNNT*s (*TNNT1-4*) (fig. 5B).

**Figure 5.**
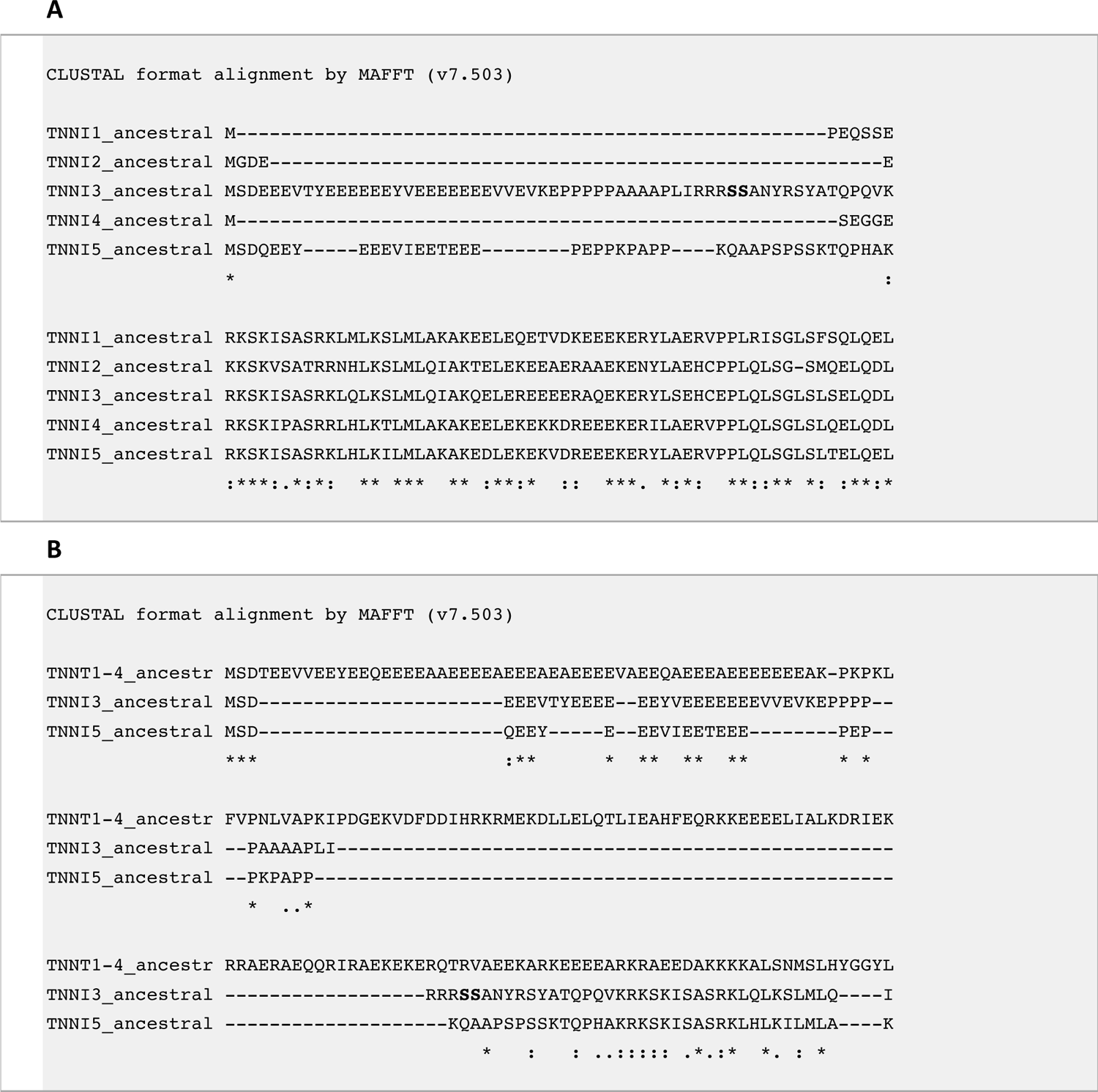
Alignments of N-terminal portions of *TNNI* and *TNNT* genes. A) Similar glutamic acid and proline rich N-terminal extensions in predicted ancestral *TNNI3* and *TNNI5*. Bolded text (serines) indicates protein kinase A target site in *TNNI3* that is absent in *TNNI5*. B) comparison of *TNNI3* and *TNNI5* with ancestral *TNNT* (common ancestral sequence of *TNNT1-4*) also shows similarities in N-terminal extensions.

### Cardiac and skeletal muscle gene and protein expression

Having established that cartilaginous and ray-finned fish *TNNI5* shares a strikingly similar N-terminal extension with tetrapod and sarcopterygian fish *TNNI3*, we next investigated the expression of different *TNNI*s in cardiac and skeletal muscle of diverse cartilaginous fishes, non-teleost ray-finned fishes, early diverging teleosts, and a sarcopterygian fish. Gene expression (analysis of previously published RNA-seq data; see Supplementary Material online for full species list and data accession information) and protein expression (Western blotting and mass spectrometry) analysis were conducted with a particular focus on the expression and characterization of the intriguing *TNNI5* paralog.

In the cardiac transcriptomes of cartilaginous fishes, we found the expression of a broad array of *TNNI*s. In small-spotted catshark (*S. canicula*), for instance, we identified transcripts for each of the four genes found in the genome, whereby *TNNI1* and *TNNI5* were dominantly expressed (∼33% *TNNI1* and ∼66% TNNI5), *TNNI4* exhibited only low expression (< 0.5% *TNNI*) and *TNNI2* was found only at trace levels (0.1% *TNNI*) (fig. 6A). In most of the other sharks (*i.e.* Great white shark (*Carcharodon carcharias*), Great hammerhead shark (*Sphyrna mokarran*), shortfin mako shark (*Surus oxyrinchus*)) as well as yellow stingray (*Urobatis jamaicensis*), we likewise found mixed expression of *TNNI1* and *TNNI5* and that the other genes were also absent or expressed at negligible levels. In Greenland shark (*Somniosus microcephalus*) *TNNI5* was strongly dominant (>90% *TNNI*). In the chimaerid elephant fish (*C. milii*), *TNNI1* was almost exclusively expressed (>97% *TNNI* expression), although *TNNI5* transcripts were also detected (accounting for the remaining 3% *TNNI*). In almost all of the ray-finned fishes, including ‘basal’ (early-diverging) actinopterygians (Senegal bichir (*Polypterus senegalus*), paddlefish (*Polyodon spathula*), spotted gar (*Lepisosteus oculatus*), bowfin (*Amia calva*)) as well as early-diverging teleosts (European eel (*Anguilla anguilla*) and silver arowana (*Osteoglossum bicirrhosum*)), *TNNI1* was the virtually exclusively expressed paralog, although in Siberian sturgeon (*Acipenser baerii*) there was also low but evident (∼3% *TNNI*) expression of *TNNI4* (fig. 6A). In African lungfish (*P. annectens*) we identified slightly predominant gene expression (55% TNNI) of *TNNI3*, with the remainder comprising of *TNNI1*.

**Figure 6.**
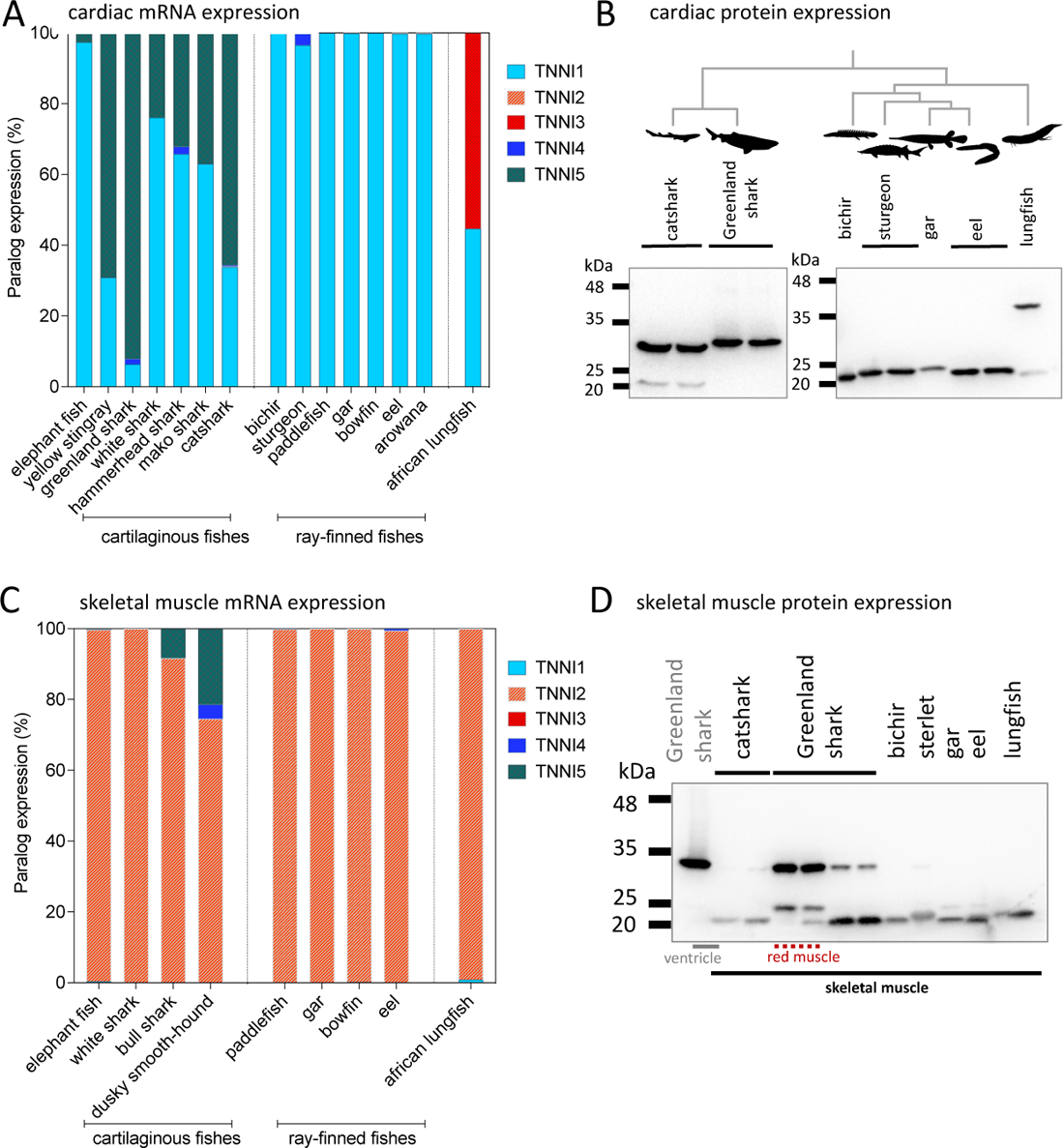
*TNNI* gene and TnI protein expression in cardiac and skeletal muscle of gnathostome vertebrates. A,C) gene expression as studied by transcriptomics. B,D) immunoblots with general TnI antibody. Fish silhouettes are courtesy of phylopic.org.

We next studied TnI protein expression in two shark species (small-spotted catshark (*S. canicula*) and Greenland shark (*S. microcephalus*)), four early-diverging ray-finned fishes (Senegal bichir (*P. senegalus*), sterlet (*A. ruthenus*), Florida gar (*Lepisosteus platyrhincus*) and European eel (*A. anguilla*)) and African lungfish (*P. annectens*) fig. 6B), and in each case the predicted protein sizes correlated well with predictions from transcriptomics. In Greenland shark, only a relatively large TnI (∼32 kDa) was detected, aligning with the N-terminal extended *TNNI5*, whereas in catshark we were also able to observe the less abundant expression of a shorter TnI protein (∼20 kDa), corresponding with the complementary expression of *TNNI1* indicated by the transcriptome (fig. 6B). Mass spectrometry for protein identification was used to confirm the protein sequence of the dominant band matched the predicted sequence for both catshark (28% coverage) and Greenland shark (65% coverage) protein predicted from *TNNI5*. In both cases, the peptide matches included a large proportion of the N-terminal extension (Supplementary fig. S1, Supplementary Material online). In each of the ray-finned fishes we observed only a band of lower molecular mass, corresponding with the N-terminal extension-absent *TNNI1* sequences predicted from transcriptomics. In lungfish, we observed expression of both a high molecular weight (the dominant band) and lower molecular weight TnI. The dominant band was confirmed as that predicted from the *TNNI3* sequence, with a long N-terminal extension, which was verified with mass spectrometry for protein identification (83% coverage).

Given that the genomes of some early-diverging actinopterygians contained an N-terminal extended TnI (*TNNI5*) that was not abundantly expressed in their hearts, we extended our survey to skeletal muscle (fig. 6C). However, none of the species’ transcriptomes that we were able to investigate (paddlefish, gar, bowfin) showed evidence of *TNNI5* expression in skeletal muscle (fig. 6C). In all of these species, the *TNNI2* paralog was strongly dominant, which indicates the preferential dissection of fast twitch muscle in skeletal muscle samples. Western blot analysis of skeletal muscle homogenates from bichir, sturgeon, gar and eel also indicated that only one or more lower molecular weight TnIs were present (fig. 6D). Surprisingly, however, some shark skeletal muscle tissues (particularly the dusky smooth-hound (*Mustelus canis*)) expressed the N-terminal extended *TNNI5* mRNA (fig. 6C), and in Greenland shark skeletal muscle, higher molecular weight TnI was confirmed to be expressed as protein (fig. 6D), which appeared (qualitatively) more abundantly expressed in red skeletal muscle than white muscle.

### Phosphorylation of cardiac-expressed TnI by PKA

In the mammal TnI3 (*TNNI3*), PKA is known to primarily target two serine residues within a canonical PKA motif (Ser-23/24)(Martin-Garrido et al. 2018). Sequence alignment indicates the PKA motif is well conserved in all species with *TNNI3*, including coelacanth and lungfish (Rasmussen et al. 2022). However, despite containing an N-terminal extension and exhibiting cardiac expression, *TNNI5* of sharks does not contain a predicted PKA phosphorylation site in the N-terminal extension (see fig. 5). To investigate if the TnI expressed in the hearts of sharks, diverse ray-finned fishes, or lungfish are targeted by PKA, we employed a phospho-PKA motif specific antibody in Western blots on cardiac homogenates before stripping the membrane and re-probing for TnI (fig. 7). Our data showed that in lungfish, a PKA phosphorylated band co-localised with the confirmed TnI location (in agreement with another recent study (Rasmussen et al. 2022)), whereas in sharks and ray-finned fish, there was no co-localisation of PKA substrates and TnI band, consistent with predictions from the protein sequences and previous findings that the TnI of ray-finned fish exhibits little phosphorylation (Gillis & Klaiman 2011; Patrick et al. 2010). The PKA-phosphorylated band in both shark species at ∼26 kDa (fig. 7) remains unidentified, but non-TnI candidates were anticipated for the non-specific PKA substrate antibody.

**Figure 7.**
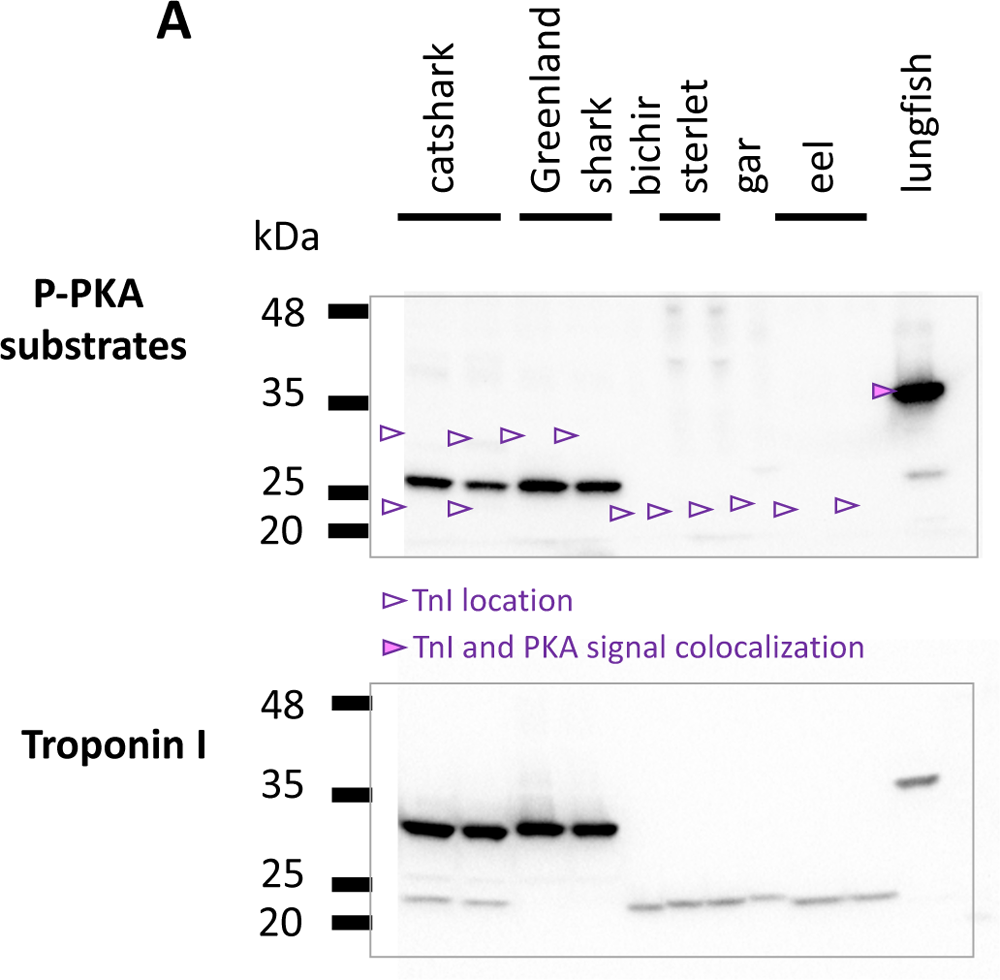
Protein kinase A-mediated phosphorylation of cardiac expressed TnI in gnathostome vertebrates. The membrane was blotted with a phospho-PKA substrate antibody, then stripped and reprobed with general TnI antibody in order to identify if a canonical PKA site was phosphorylated in TnI. Arrows show localisation of TnI on P-PKA blot. Filled arrows show co-localisation of TnI and PKA band, non-filled arrows indicate no co-localisation.

## Discussion

In this study, we combined bioinformatic searches for phylogenetic and synteny analyses with gene and protein expression studies to reconstruct the evolution of the vertebrate *TNNI* and *TNNT* gene families. Our analyses suggest a novel hypothesis regarding the duplicative history of these gene families, identify additional paralogs that are absent from mammalian genomes, and provide a strong hint that the presence of an N-terminal extension in mammalian cardiac TnI represents the retention of an ancestral state. A summary of our interpretation of *TNNI* and *TNNT* gene origins and losses is presented in fig. 8. The lineage-specific gene losses resulted in different paralog repertoires being available for tissue-dependent expression different vertebrate groups. This is well illustrated in the context of cardiac *TNNI* expression, where tetrapods and lobe-finned fishes characteristically express *TNNI3*, ray-finned fishes by and large express *TNNI1*, and cartilaginous fishes express a variable combination of *TNNI1* and *TNNI5* in the heart.

**Figure 8.**
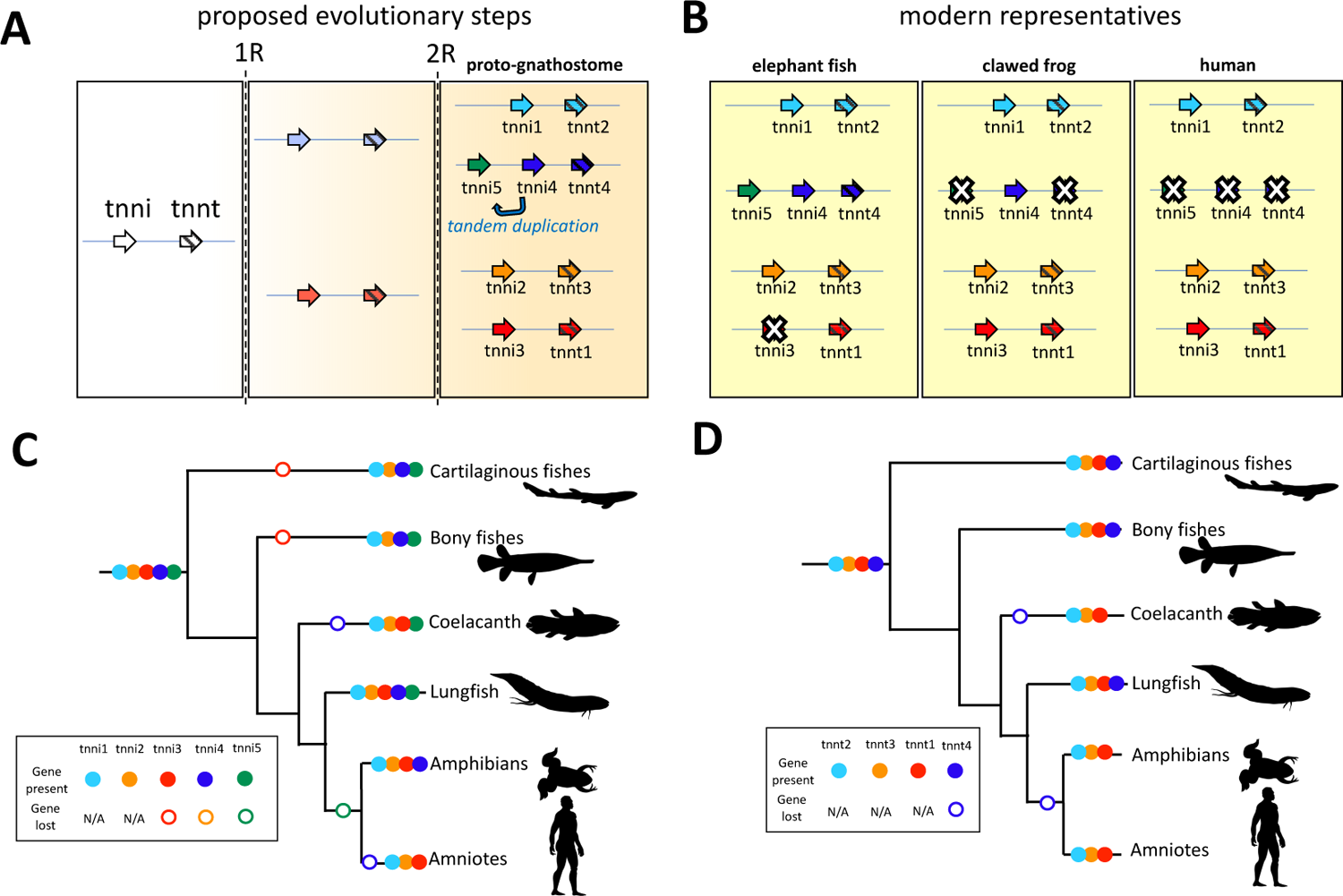
The origin and losses of *TNNI* and *TNNT* genes in vertebrates. A) predicted evolutionary steps in evolution showing how whole genome duplications (1R and 2R) and tandem duplications generated gene diversity. White crosses (X) mark genes that have been lost in extant lineages. Whilst our maximum likelihood tree suggests the tandem duplication giving rise to *TNNI4* and *TNNI5* occurred before the genome duplication that gave rise to *TNNI1* and *TNNI5*, which would require subsequent gene loss of a paralog in both gnathostomes and cyclostomes independently, it is also possible that the tandem duplication occurred only in *TNNI4* and *TNNI5* after 2R (here shown by arrow marked ‘tandem duplication’). B) Comparison of which duplicated genes were retained or lost in three specific gnathostomes (elephant fish, *C. milii,* tropical clawed frog*, X. tropicalis* and human*, H. sapiens*. C,D) broader overview of patterns of *TNNI* and *TNNT* gene loss in major vertebrate lineages. Animal silhouettes are courtesy of phylopic.org.

### TNNI and TNNT paralogs originated in 2R whole-genome duplication events

Earlier convention has been to pigeon-hole the *TNNI* paralogs found in non-mammalian vertebrates into the three classes of *TNNI* defined in mammals. By contrast here, tree reconciliation allows us to trace the five different *TNNI*s and the four different *TNNT*s of gnathostomes back to the last common ancestor of the group. Our trees also suggest that at least three of the *TNNI*s and four of *TNNT* cyclostome paralogs predate the split between hagfish and lamprey, but support for the corresponding nodes is low to move beyond this broad statement. Going deeper into the phylogeny, our maximum likelihood tree placed gnathostome *TNNI4* as sister to a cyclostome *TNNI* gene, implying that the tandem duplication giving rise to *TNNI4* and *TNNI5* would have predated the duplication that gave rise to *TNNI5* and *TNNI1* and the split between cyclostomes and gnathostomes. This would require multiple losses of *TNNI* and *TNNT* genes in both cyclostomes and gnathostomes to account for the extant repertoires. However, a tree where *TNNI4* is constrained to be sister to *TNNI5* (Supplementary fig. S2, Supplementary Material online) is not significantly different from the unconstrained tree (Supplementary Table 1, Supplementary Material online). This constrained tree requires fewer gene losses and maps the tandem duplication giving rise to *TNNI4*-*TNNI5* to the last common ancestor of gnathostomes, which is consistent with the observed phyletic distribution of the genes. Thus, our analyses indicate the presence of four different *TNNI*-*TNNT* pairs in the last common ancestor of gnathostomes, and of at least three *TNNI*-*TNNT* clusters in the last common ancestor of cyclostomes.

Unfortunately, our phylogenies lack power to resolve relationships between gnathostome and cyclostome *TNNI*s and *TNNT*s and move ancestral reconstruction deeper. This is not surprising because cyclostome genomes are unusual with respect to nucleotide, codon and amino acid composition (Qiu et al. 2011; Kuraku 2013) that complicate the resolution of orthology based on phylogenies. In some cases, synteny is informative to resolve ambiguous gene phylogenies (Kuraku & Meyer 2012; Hoffmann et al. 2010; Campanini et al. 2015), but in others such as the *TNNI*-*TNNT* pairs, synteny shows similarities with gnathostomes but ultimately is not informative, as in the globin X genes of vertebrates (Hoffmann et al. 2021). Nevertheless, the presence of multiple *TNNI*-*TNNT* pairs in both groups provides a strong indication that the last common ancestor of vertebrates possessed four *TNNI*-*TNNT* pairs in its genome.

Whole-genome duplications played a critical role in expanding the repertoire of vertebrate genes. The presence of four different *TNNI-TNNT* clusters in gnathostomes which are phylogenetically arranged by location and are flanked by additional gene families that appear to have co-duplicated with the *TNNI-TNNT* suggest that the *TNNI-TNNT* clusters of vertebrates also expanded via WGDs, a notion previously speculated on more limited evidence (Shaffer & Gillis 2010). The presence of independent duplications in cyclostomes and synteny similarities within the cyclostome clusters are also consistent with this interpretation. Further, the three human *TNNI-TNNT* pairs and the three lamprey *TNNI-TNNT* pairs all map to proto-vertebrate chromosome Pv11 from Nakatani et al. (2021), as does the tandem of *TNNT* genes in amphioxus (Supplementary Table 2, Supplementary Material online). All of these observations suggest that the vertebrate *TNNI-TNNT* gene families expanded as a result of WGDs.

There are competing hypotheses regarding the number and timing of the WGDs early in vertebrate evolution. There is consensus that gnathostomes underwent two rounds of WGD (Meyer & Schartl 1999; McLysaght et al. 2002; Dehal & Boore 2005), 1R and 2R, and that 1R predates the split of extant cyclostomes and gnathostomes. The placement of 2R on the vertebrate tree, however, is controversial (Kuraku et al. 2009). Whereas some authors place 2R in the common ancestor of cyclostomes and gnathostomes (Sacerdot et al. 2018), *i.e.* ‘2R-early’, more recent studies place 2R in the last common ancestor of gnathostomes (Simakov et al. 2020; Nakatani et al. 2021), *i.e.* ‘2R-late’, and suggest that cyclostomes underwent an independent polyploidization early in their evolution (Mehta et al. 2013; Nakatani et al. 2021). These competing explanations make alternative phylogenetic predictions (Supplementary fig. S3. Supplementary Material online). Our results do not fit either the 2R-early or 2R-late hypotheses in strict sense but they are easier to reconcile with the 2R-late hypothesis. Linking the duplicative history to the 1R and 2R WGDs is trivial in the case of gnathostomes, the *TNNI2/3-TNNT1/3* and *TNNI1/4/5-TNNT2/4* pro-orthologs would derive from 1R, which then expanded to the four different *TNNI-TNNT* pairs we see today, with the *TNNI4/5* pro-ortholog in single copy state. The case of cyclostomes is more complex. The *TNNI-TNNT* clusters of lamprey share synteny similarities that distinguish them from the *TNNI-TNNT* clusters of gnathostomes. This fits well with the 2R-late hypothesis, which posits that cyclostomes underwent an independent polyploidization event. Under this scenario, the *TNNI116956477*-*TNNT116956460* and *TNNI116939854*-*TNNT116939851* pairs of lamprey and the *TNNI2*-*TNNT3* and *TNNI3*-*TNNT1* pairs of gnathostomes would represent independent expansions of one of the post 1R *TNNI*-*TNNT* pairs, and whereas the *TNNI1-TNNT2* and *TNNI4/5-TNNT4* pairs of gnathostomes and the *TNNI116945613-TNNT116945609* would derive from the other post-1R *TNNI-TNNT* pair. We favour this interpretation because it requires less changes relative to the observed trees and the synteny similarities within cyclostomes and within gnathostomes. Unfortunately, support for the relevant nodes is low, and topology tests are not informative (Supplementary Table 1, Supplementary Material online).

### Relationships among the TNNI and TNNT paralogs

In contrast to previous studies, which indicated that *TNNI1* and *TNNI3* were more closely related to one another than to *TNNI2* (Hastings 1997; Shaffer & Gillis 2010; Sheng & Jin 2016), our expanded analyses place *TNNI3* as sister to *TNNI2*, and place *TNNI1* as sister to *TNNI5*, with *TNNI4* grouping with the *TNNI1+TNNI5* clade. This has important implications regarding the origin of the TNNI3 terminal extension (see below). The parallel analyses of *TNNI* and *TNNT*, consistently found in close proximity in vertebrate genomes, provided a powerful tool to cross-examine predicted gene duplications. In support of the surprising *TNNI2*-*TNNI3* sister relationship, we also found a sister relationship between their syntenically associated *TNNT* genes, *TNNT3* and *TNNT1*, respectively. Likewise, the *TNNT2*-*TNNT4* affinity supported the close relationship of *TNNI1*, *TNNI4* and *TNNI5*.

### Origin of the troponin complex

The three subunits of troponin, *TNNC*, *TNNI*, and *TNNT* can be traced back to at least to the last common ancestor of bilateria (Barnes et al. 2016; Yaguchi et al. 2017). Our synteny analyses provides some hints as to how this three-subunit complex emerged. *TNNI* and *TNNT* are the most similar subunits to one another, which probably emerged as tandem duplicates deep in the past (Chong & Jin 2009). To the best of our knowledge (cf. Herranz et al. 2005), vertebrates are the only group that have retained this close tandem arrangement. Interestingly, in amphioxus, an invertebrate chordate, *TNNT* and *TNNC* are in a cluster. We speculate that the three proteins could potentially have been in a single cluster in the early stages of their evolution, and that amphioxus might reflect the ancestral condition of a *TNNC* gene in cluster with the progenitor of *TNNI* and *TNNT*. The original troponin subunit might have been made of a *TNNC* chain associated with two chains contributed by this *TNNI*/*T* pro-ortholog. In time, a tandem duplication gave rise to *TNNI* and *TNNT* and a sub functionalization process allowed each of these subunits to evolve the more specific roles they have today. In the vertebrate tree of life, the emergence of multiple *TNNI* and *TNNT* subunits via WGD allowed further specialisations of these different genes, which acquired more refined muscle tissue specificity in this group of animals. Such specialisation enabled muscle type specific functional phenotypes.

### *TNNI1* is expressed in the fish heart

Virtually all of the previous phylogenetic studies on vertebrate *TNNI* evolution (Sheng & Jin 2016; Shaffer & Gillis 2010; Gross & Lehman 2016; Rasmussen et al. 2022) have shown that the teleost cardiac-expressed *TNNI* gene clusters within *TNNI1* (ssTnI) of tetrapods, yet it is still frequently labelled as a fish ‘cTnI’ which implies orthology with *TNNI3.* Its phylogenetic ‘misplacement’ has been repeatedly attributed to its lack of N-terminal extension (Shaffer & Gillis 2010; Rasmussen et al. 2022). By combining our phylogenetic analysis with comparisons of conserved synteny and a broad transcriptomic survey, we unequivocally conclude that ray-finned fishes, including teleosts, lack the *TNNI3* gene and simply express a *TNNI1* ortholog in the heart. This is consistent with the state in embryonic and neonatal mammals, which express *TNNI1* in the heart before *TNNI3* becomes exclusively expressed as juveniles and adults (Saggin et al. 1989; Reiser et al. 1994). In the adult mammalian heart, overexpression of ssTnI (*TNNI1*) at the expense of cTnI (*TNNI3*) confers increased tolerance to acidosis (Wolska et al. 2001), and may provide similar benefits in the fish heart, which in many species show exceptional performance during acidosis (Driedzic & Gesser 1994; Hanson et al. 2009; Joyce et al. 2015).

Given that ray-finned fishes have lost the *TNNI3* gene (which encodes for cTnI in mammals), we suggest that the use of the ‘cTnI’ name for cardiac expressed *TNNI*s in these fish, as has been frequently applied (Sheng & Jin 2016; Shaffer & Gillis 2010; Alderman et al. 2012; Gillis & Klaiman 2011), should be discontinued. Indeed, more generally the currently used protein nomenclature is based on similarities to human genes, some of which are absent from vertebrate genomes, and incorporate information about the tissue where the protein is found and does not align well with our evolutionary hypothesis. Because non-mammalian species, such as teleost fish (Alderman et al. 2012; Shih et al. 2015), have multiple *TNNI* genes in different striated muscle types, and non-orthologous genes may be expressed in a given muscle type, it becomes ambiguous to use protein nomenclature based on muscle type specific expression of mammals. We therefore advocate that protein names be derived from the gene number (*i.e.* TnI1-5) in studies that include non-mammalian vertebrates.

Even in some cartilaginous fishes and lungfish, *TNNI1* was relatively highly expressed in the heart, but unlike in ray-finned fishes it was found in combination with another paralog, *i.e. TNNI5* or *TNNI3* respectively. The ability to express two or more distinct *TNNIs* with different properties (*i.e. TNNI* multiplicity) may provide a substrate for acclimation to different environmental conditions (Alderman et al. 2012). Such an ability would be of obvious benefit for ectothermic species and has also been demonstrated to occur with respect to TnC paralog expression in the trout heart with thermal acclimation (Genge et al. 2013).

### A common origin for the N-terminal extension in vertebrate *TNNI*

The N-terminal extension peptide in TnI3 is widely viewed as an evolutionary novelty that appeared in the sarcopterygian fish and tetrapod lineage (Sheng & Jin 2016; Palpant et al. 2010; Shaffer & Gillis 2010; Rasmussen et al. 2022). However, we identified that the *TNNI5* in cartilaginous fishes, non-teleost ray-finned fishes, and coelacanth contained an N-terminal extension with striking similarity to that found in *TNNI3*, particularly of lungfish and amphibians.

The sister relationships of *TNNI3* with *TNNI2*, and *TNNI5* with *TNNI1* and *TNNI4*, were robustly supported (and cross-supported by the tree of syntenic *TNNT*s), indicating that *TNNI3* and *TNNI5* are only distantly related. Based on the molecular similarity of the N-terminal extensions in the proteins encoded by *TNNI5* and *TNNI3*, it appears likely that it was found in the common ancestor of vertebrate *TNNI*s and was independently lost in *TNNI1*, *TNNI2* and *TNNI4* lineages. The common origin of the N-terminal extension is also supported by the N-terminal extension in the *TNNI* of *C. intestinalis*, which is structurally similar to *TNNI3* of tetrapods such as the tropical clawed frog (MacLean et al. 1997). This earlier led Hasting’s to also conclude that the N-terminal extension could be ancestral (Hastings 1997), although this hypothesis has largely been overlooked in more recent work (Sheng & Jin 2016; Palpant et al. 2010; Shaffer & Gillis 2010; Rasmussen et al. 2022). Some protostome TnI genes also contain an N-terminal extension, and the possibility that it was conserved with vertebrate *TNNI3* has also been previously acknowledged (Cao et al. 2019; Barnes et al. 2016). Indeed, *TNNT*, as the sister family to *TNNI* (Chong & Jin 2009) that diverged following a duplication before the separation of protostomes and deuterostomes (Cao et al. 2019), also contains a proline- and glutamic acid-rich N-terminal extension. An alignment of the ancestral vertebrate *TNNT* (prior to 2R) with ancestral *TNNI3* and *TNNI5* reveals stretches of similarity with TnT in the N-terminus (fig. 5B), indicating the *TNNI* N-terminal extension may date back to before the *TNNI-TNNT* separation.

That the N-terminal extension was lost multiple times in other vertebrate *TNNI* lineages is initially surprising but is also supported by evidence that the single *TNNI* gene of *C. intesintalis* is alternatively spliced, with the N-terminal extension expressed only in cardiac muscle but excluded in skeletal muscle (MacLean et al. 1997). This indicates that it may provide a benefit to lose the N-terminal extension in skeletal muscle, which was only afforded at the genomic level following the gene duplications that generated paralog diversity (Hastings 1997; MacLean et al. 1997).

In the mammalian heart, TnI3 is a major target for PKA following activation by ß-adrenergic stimulation (Bers et al. 2019) where it affects myofilament Ca^2+^ sensitivity (Fentzke et al. 1999; Robertson et al. 1982). Whilst *TNNI3,* in sarcopterygian fishes and tetrapods, and *TNNI5,* in sharks and rays, are both abundantly expressed in the heart, an important distinction is that TnI5 appears to lack functional PKA phosphorylation target sites in the N-terminal extension. This would presumably limit sensitivity of cardiac myofilaments to the effects of adrenergic stimulation (Pi et al. 2002) and reduce the functional scope of the heart. *TNNI5* also appears to be expressed in shark skeletal muscle (red muscle in particular), whereas *TNNI3* is strictly only found in the heart in mammals (Sheng & Jin 2016). Given its unique structure and expression pattern, it would be of interest for future work to establish the functional properties of cartilaginous fish TnI5, including the possible influence of its non-phosphorylatable N-terminal extension.

## Conclusion

Our analyses suggest a novel hypothesis regarding the expansion of the *TNNI* and *TNNT* gene families of vertebrate, linking the presence of multiple *TNNI-TNNT* pairs in their genomes to the WGDs early in the history of the group. Our results show how a combination of tandem gene duplications and whole-genome duplications have worked together to generate protein diversity that allow the differentiation of different muscle types of troponins. Under the 2R-late hypothesis, our analyses suggest that the presence of four *TNNI-TNNT* clusters in the genomes of gnathostomes are the product of the 1R and 2R WGDs. In cyclostomes, which share 1R with gnathostomes, the presence of multiple *TNNI-TNNT* pairs seems to be a combination of 1R and a polyploidization event specific to this lineage. Moving closer to present, we also identify additional paralogs present in the last common ancestor of gnathostomes that are absent from mammalian genomes. The genes were retained by a subset of extant lineages, such as the *TNNI3* from amniotes that is absent in cartilaginous fish or ray-finned fishes, or the *TNNI4/5-TNNT4* locus, which has apparently been lost in amniotes. Moving deeper in time, our results potentially suggest that the *TNNC-TNNI-TNNT* genes could have been arranged in a cluster in the common ancestor of bilaterians, estimated to have lived ∼850-700 million years before present. Our new evolutionary framework highlights the need for revised nomenclature to more faithfully portray the evolutionary affiliations of some previously mis-annotated genes (see Table 1), and we also provide consistent names for previously unnamed *TNNI* genes (*e.g.* ‘slow skeletal-like’ troponin I genes found across cartilaginous and ray-finned fish lineages that can now be identified as *TNNI4* or *TNNI5*).

We found that two distantly related lineages, *TNNI3* and *TNNI5*, encode TnI proteins with remarkably similar N-terminal extensions (fig. 5), which is most easily explained by it being present in their common ancestor and independently lost in *TNNI1*, *TNNI2* and *TNNI4* lineages. The discovery of a ‘second’ vertebrate *TNNI* with an N-terminal extension provides the strongest evidence to date that the extension, which has been widely viewed as unique to *TNNI3*, could represent an ancestral state of gnathostome TnI prior to the 2R duplication events, and likely dates back even further to the origin of bilaterian *TNNI*. Shark hearts exhibited dominant protein expression of the N-terminal extended *TNNI5*. However, the heavily studied PKA-target phosphorylation sites present in mammal TnI3 were only found in the *TNNI*3 gene family and not *TNNI5*.

## Materials and Methods

### Bioinformatic searches and curation

Our strategy was to include all known *TNNI* and *TNNT* genes in a broad range of vertebrates with annotated whole genome assemblies. We pre-defined the following target species: three distantly related species of cartilaginous fish (elephant fish, *Callorhinchus milii,* thorny skate, *Amblyraja radiata,* and small-spotted catshark, *Scyliorhinus canicula*), three non-teleost ray finned fishes (reedfish, *Erpetoichthys calabaricus,* sterlet sturgeon, *Acipenser ruthenus* and spotted gar, *Lepisosteus oculatus*), two distantly-related teleosts (Asian bonytongue, *Scleropages formosus* and zebrafish, *Danio rerio*), African coelacanth (*Latimeria chalumnae*), West African lungfish (*Protopterus annectens)*, an amphibian (tropical clawed frog, *Xenopus tropicalis*), a non-avian reptile (anole lizard, *Anolis carolinensis*), a bird (chicken, *Gallus gallus*), a monotreme (Australian echidna, *Tachyglossus aculeatus*) and a eutherian mammal (human, *Homo sapiens*). We additionally included two cyclostomes, the sea lamprey (*Petromyzon marinus*) and inshore hagfish (*Eptatretus burger*). An amphioxus (Florida lancelet, *Branchiostoma floridae*) and a tunicate (vase tunicate, *Ciona intestinalis)* were included as outgroups. We used both National Center for Biotechnology Information (NCBI) and Ensembl (Release 105) databases. The NCBI database gene pages were manually searched (*i.e.* “species name + “troponin I”) and also searched with protein-protein BLAST using known (*i.e.* mammalian or tropical clawed frog) TnI sequences, and *TNNI* genes were searched for a given species in Ensembl. Where genes of a given species appeared on both NCBI and Ensembl, the former was typically used. The African lungfish genome has only recently been sequenced (Wang et al. 2021) and predicted proteins from the gene annotations appeared inconsistent with other species. As we assembled transcriptomes for African lungfish cardiac and skeletal muscle (below), which generated more plausible and complete *TNNI1*, *TNNI2* and *TNNI3* sequences, these were used instead for the phylogenetic analyses. The transcriptome-predicted lungfish *TNNI3* is consistent with that recently cloned by Rasmussen et al. (2022) in the same species. One annotated amphioxus *TNNT* gene (NCBI 118422967) was excluded as it shared little resemblance with *TNNT* in any other species.

### Phylogenetic analysis

*TNNI* and *TNNT* alignments were generated using MAFFT (v7.490) using the einsi and linsi strategies (Katoh & Standley 2013). The resulting alignments were compared using MUMSA (Lassmann & Sonnhammer 2006) and the alignments with the highest scores were selected for downstream processing. Phylogenetic relationships were estimated using IQ-TREE (multicore version 2.2.0-beta) (Minh et al. 2020). First, the best-fitting model of amino acid substitution was selected using the ModelFinder subroutine from IQ-Tree (Kalyaanamoorthy et al. 2017; Katoh & Standley 2013). Then, searches were run under the selected model using 10,000 pseudoreplicates of the ultrafast bootstrap procedure to assess support for the nodes (Hoang et al. 2018). Competing phylogenetic hypotheses were compared using the approximately unbiased test (Shimodaira 2002) as implemented in IQ-Tree. The alignments, tree files, and a log of the commands required to replicate our results are all available as a compressed file with the Supplementary Material online.

### Synteny

Synteny diagrams were generated for key species (amphioxus, tunicate, lamprey, elephant fish, catshark, spotted gar, zebrafish, coelacanth, tropical clawed frog, human) for genomic regions harbouring select *TNNI*/*TNNT* genes using the ‘Genomic context’ section on the relevant NCBI gene pages.

### Reconstruction of ancestral protein sequences

Common ancestral protein sequences for each TNNI paralog and the common vertebrate TNNT were predicted using FireProt ASR (ancestral sequence reconstruction) v1.1 web server using default parameter settings (Khan et al. 2021). An unrooted tree was generated with the same alignments as used for our phylogenetic trees, except that for TNNT the “bonytongue_tnnt2a” (XP_029102145.1) sequence was omitted as it contained an unknown amino acid (‘X’, *i.e.* low quality prediction) so was not recognised by the software. Human *TNNI3* and *TNNT1* were used as ‘query’ sequences. Nodal sequences from the common ancestor of *TNNI1, TNNI2, TNNI3, TNNI4* and *TNNT5*, and the common ancestor of vertebrate *TNNT1-4* were exported and aligned using MAFFT (v. 7.503).

### Gene expression

Previously generated heart and skeletal muscle RNA-seq data for a broad cohort of cartilaginous fishes, ray-finned fishes and lungfish were collated (see Supplementary Material online for full list of species and SRA accession numbers). Sequences with a low-quality score (regions averaging a score <5 over a 4bp sliding window, and leading/trailing sequences scoring <5) were removed using Trimmomatic (Bolger et al. 2014). The cleaned reads were processed in Trinity 2.8.4 using default parameters (Haas et al. 2013; Grabherr et al. 2011). Open reading frames (ORFs) were predicted from the transcripts using Transdecoder (https://github.com/TransDecoder/TransDecoder/wiki) with a minimum length threshold of 100 amino acids. CD-HIT (Li & Godzik 2006) was used to eliminate redundancy by clustering nucleotide sequences with ≥99% similarity. The reads were mapped using Bowtie-2 (Langmead & Salzberg 2012), and pseudo-aligned to the predicted ORFs using Kallisto (Bray et al. 2016), allowing relative abundance estimates of each transcript to be calculated. Annotation was performed using BLAST (blastp, default cut-offs) searches to query a broad range of *TNNI* genes (*TNNI1,2,4,5* from catshark and *TNNI3* from tropical clawed frog). The top hits were manually curated to find *TNNI* paralogs by blastp (default settings) against the NCBI database and candidate genes were cross checked against our conserved synteny diagrams to confirm their identity.

### Protein expression and phosphorylation immunoblots

Small-spotted catshark (*Scyliorhinus canicula*; 600-800 g; N=2), Senegal bichir (*Polypterus senegalus*; ∼15 g, N=2), sterlet (*Acipenser ruthenus*; ∼40 g N=2), Florida gar (*Lepisosteus platyrhincus*: ∼ 2 g, N=2), European eel (*Anguilla anguilla*; 600-800 g; N=2), and West African lungfish (*Protopterus annectens*; 410 g; N=1) were obtained from local commercial dealers and euthanised with an overdose (1 g L^-1^) of bicarbonate-buffered tricaine methanesulfonate (MS-222) followed by destruction of the brain, a procedure endorsed by the local animal experiments committee and in accordance with Schedule 1 of the Home Office Animals (Scientific Procedures) Act 1986. The heart (ventricle) and skeletal muscle (epaxial muscle) were dissected and rapidly frozen on dry ice and stored at −80°C. Samples from non-sexually mature Greenland sharks (*Somniosus microcephalus*; N = 2, TL=398 and 386 cm) were collected in 2021 in south-eastern Greenland from the Danish research vessel Dana. Sharks were caught via long lines at depths between 285-325 m. Immediately after capture and euthanisation, samples of ventricle, and white skeletal muscle and red skeletal muscle, were flash frozen in liquid nitrogen and stored at −80 °C until use.

All experiments were conducted with tissue stored (at −80°C) for less than 6 months, as initial pilot experiments indicated that catshark tissue that was several years old showed greatly reduced general phosphorylation levels. Tissue samples were homogenized in 10 µl mg^-1^ RIPA buffer (Millipore, 20-188) with 1% protease and phosphatase inhibitor (PPI) cocktail (PPC1010 Sigma-Aldrich). Protein concentration was measured with bicinchoninic acid (BCA) assay (71285-3 EMD Millipore) before samples were denatured with 2x Laemmli buffer (S3401 Sigma-Aldrich) and boiled at 95°C for 5 min. Tris-glycine gels (10 or 12.5% acrylamide) were run with an Invitrogen XCell SureLock Mini-Cell and transferred to a PVDF membrane with XCell II Blot Module (ThermoFisher). For the specific gel used for each blot, alongside raw uncropped blot images, see Supplementary Material online. Gels were run at 200 V for 45 min. 15 µg protein was loaded in each lane and run alongside 5 µL BLUeye Prestained Protein Ladder (Sigma-Aldrich 94964). Blots were blocked for 1 h at room temperature using 5% skimmed milk in standard tris-buffered saline with 0.05% Tween 20 (TBS-T).

For the initial determination of TnI molecular weight, blots were incubated with a mouse monoclonal primary antibody against troponin I (C-4) (Santa-Cruz SC-133117). This antibody, which recognised the majority of the TnI protein except for the ‘cTnI-specific’ N-terminal extension, is recommended by the manufacturer for detection of diverse (cardiac and skeletal) TnI proteins. 1:1000 dilutions of the 200 µg ml^-1^ stock were diluted in 2% milk TBS-T to incubate the block overnight on a shaker at 4°C. The next day, a corresponding HRP-conjugated anti-mouse secondary antibody (Santa-Cruz sc-516102) at 1:5000 dilution of 400 µg ml^-1^ stock in 2% milk TBS-T was used for a 1 h incubation at room temperature. Blots were imaged on a Bio-Rad ChemiDoc and chemiluminescent signals developed with Millipore/Immobilon Classico Western HRP substrate (Merck WBLUC0500) or Bio-Rad Clarity Western ECL Substrate (Bio-Rad 1705061).

To identify phosphorylated protein kinase A (phosho-PKA) substrates (RRXS*/T*) in heart samples, we used the New England Biolabs 9624S rabbit primary antibody (1:1000 dilution of unspecified stock concentration) at 4°C overnight with gentle agitation followed by goat anti-rabbit secondary antibody (New England Biolabs 7074P2; 1:3000 dilution of unspecified stock concentration) for 1 h at room temperature. The blot was imaged as above, then stripped (Restore Western Blot Stripping Buffer, ThermoFisher 21059) and the TnI antibody protocol followed as described above.

### Mass spectrometry

To verify if the apparently N-terminal extended dominant TnI proteins present in catshark and Greenland shark as well as African lungfish corresponded with the predicted *TNNI5* or *TNNI3* sequences, protein identification with liquid chromatography-mass spectrometry (LC-MS) was performed with the University of Manchester Bio-MS Research Core Facility (RRID SCR_020987). 20 µg protein per lane was run on a 16% acrylamide Tris-glycine gel (Invitrogen XP00165BOX) which was then stained with SimplyBlue™ SafeStain (ThermoFisher LC6065). The band at the location corresponding with TnI identified by the immunoblot was excised and digested with elastase. The samples were analysed with LC-MS/MS using an UltiMate 3000 Rapid Separation LC (RSLC, Dionex Corporation, Sunnyvale, CA) coupled to an Orbitrap Exploris 480 (Thermo Fisher Scientific, Waltham, MA) mass spectrometer. Mobile phase A was 0.1% formic acid in water and mobile phase B was 0.1% formic acid in acetonitrile. The products were analysed with Scaffold 5 (Proteome Software, Portland, OR, USA) and searched against an in-house database including the transcriptomics-predicted *TNNI3* and *TNNI5* sequences from each respective species.

## Acknowledgements

This work was supported by Novo Nordisk Foundation (NNF19OC0055842) to WJ. Professors John Steffensen, Peter Bushnell, Diego Bernal, and Eurofleets+, and scientists and crew of the RV Dana are thanked for providing the Greenland shark heart and muscle samples.

## Data Availability Statement

The data underlying this article are available in the article and in its online supplementary material.

