## Supplemental Table S1 for "A revised perspective on the evolution of troponin I and troponin T in vertebrates"

**Supplementary Table 1.** Statistical comparisons of constrained trees.

TNNI

| Tree | logL | deltaL | bp-RELL | p-KH | p-SH | c-ELW | p-AU |
|---|---|---|---|---|---|---|---|
| 1 | -14437.73 | 0 | 0.553 | 0.776 | 1 | 0.534 | 0.768 |
| 2 | -14442.38 | 4.7 | 0.301 | 0.32 | 0.356 | 0.297 | 0.336 |
| 3 | -14439.80 | 2.1 | 0.146 | 0.224 | 0.551 | 0.169 | 0.233 |

Tree 1: Unconstrained maximum likelihood tree (figure

Tree 2: Gnathostome TNNI4 constrained as sister to Gnathostome TNNI5

Tree 3: Constrained to match the 2R-late hypothesis.

TNNT

| Tree | logL | deltaL | bp-RELL | p-KH | p-SH | c-ELW | p-AU |
|---|---|---|---|---|---|---|---|
| 1 | -19442.77 | 0 | 0.896 | 0.896 | 1 | 0.895 | 0.904 |
| 2 | -19467.96 | 25.191 | 0.104 | 0.104 | 0.104 | 0.105 | 0.0958 |

Tree 1: Unconstrained maximum likelihood tree (figure

Tree 2: Constrained to match the 2R-late hypothesis.

deltaL : logL difference from the maximal logl in the set.

bp-RELL : bootstrap proportion using RELL method (Kishino et al. 1990).

p-KH : p-value of one sided Kishino-Hasegawa test (1989).

p-SH : p-value of Shimodaira-Hasegawa test (2000).

c-ELW : Expected Likelihood Weight (Strimmer & Rambaut 2002).

p-AU : p-value of approximately unbiased (AU) test (Shimodaira, 2002).
