## Supplemental Table S2 for "A revised perspective on the evolution of troponin I and troponin T in vertebrates"

**Supplementary Table 2**

Tracing of human, sea lamprey and amphioxus genes to the proto-gnathostome, proto-cyclostome and proto-vertebrate chromosomes defined by Nakatani et al. (2021)

Human

| Gene name | NCBI Gene ID | NCBI Protein ID | Ensembl Gene ID | Ensembl Protein ID | Genomic segment | Chromosome | Proto-gnathostome Chromosome | Proto-vertebrate Chromosome |
|---|---|---|---|---|---|---|---|---|
| TNNI1 | 7135 | NP_003272 | ENSG00000159173 | ENSP00000354488 | Human_10 | 1 | 24 | Pv11 |
| TNNI2 | 7136 | NP_003273 | ENSG00000130598 | ENSP00000371331 | Human_81 | 11 | 25 | Pv11 |
| TNNI3 | 7137 | NP_000354 | ENSG00000129991 | ENSP00000341838 | Human_130 | 19 | 27 | Pv11 |

Sea lamprey

| Gene Name | NCBI Gene ID | kPetMar1 ID | kPetMar1 chr. | gPMar100 ID | gPMar100 scaffold | Lamprey chromosome fragment | proto cyclostome | proto vertebrate chromosome |
|---|---|---|---|---|---|---|---|---|
| TNNI | 116939854 | PMZ_0053578-RA | 7 | PMZ_0008467-RA | scaf_00014 | Sea Lamprey_44 | Pcc11A | Pv11 |
| | 116939854 | PMZ_0053578-RA | | (PMZ_0047782-RA) | | | | |
| TNNI | 116956477 | PMZ_0061710-RA | 65 | PMZ_0033355-RA | scaf_00064 | Sea Lamprey_106 | Pcc11E | Pv11 |
| TNNI | 116945613 | PMZ_0058172-RA | 24 | PMZ_0040924-RA | scaf_00002 | Sea Lamprey_15 | Pcc11B | Pv11 |

Amphioxus

| Gene | Gene ID | Protein ID Bfl_VNyyK (GCA_000003815.2) | Chromosome Bfl_VNyyK | Protein ID Version 2 (GCF_000003815) | scaffold version 2 | proto vertebrate chromosome |
|---|---|---|---|---|---|---|
| TNNT | 118413375 | XP_035672621 | 4 | missing in annotation | scaffold 230 | Pv11 |
| TNNT | 118414897 | XP_035675082 | 4 | XP_002590197 | scaffold 230 | Pv11 |
