## Supplemental Figures S1-S3 for "A revised perspective on the evolution of troponin I and troponin T in vertebrates"

>catsharkTNNI5
MSDQEEYEEEVVEEMEETEEVIESEPEPPKPAPPPKAAPAPRGPGAQKETIAKKSC
KISASRKLHLKILMLGKAKEDLEKETVDRNEEKEKFLAERVPPLNFTGLSLTDLQNLC
KELHQRIEIVDEERYDFEFKAGKNNYEIHDLSLKILDLRGKFKRPTLRRVRVSADAML
RALLGSKHKVSMDLRANLKSVKKDDTEKERNVEVSDWRKNVEAKSGMEGRKKMF
DAANQ

**cu|M00501|csSSX, 26,514.1 Da**
**csSSX (William Joyce (Shiels))**
**7 exclusive unique peptides, 7 exclusive unique spectra, 18 total spectra, 65/231 amino acids (28% coverage)**

MSDQEEYEEE VVEEMEETEE VIESEPEPPK PAPPPKAAPA PRGPGAQKET IAKKSCKISA
SRKLHLKILM LGKAKEDLEK ETVDRNEEKE KFLAERVPPL NFTGLSLTDL QNLCKELHQR
IEIVDEERYD FEFKAGKNNY EIHDLSLKIL DLRGKFKRPT LRRVRVSADA MLRALLGSKH
KVSMDLRANL KSVKKDDTEK ERNVEVSDWR KNVEAKSGME GRKKMFDAAN Q

>greenlandsharkTNNI5
MSDQEEQYEEEVIDETEETKEVIEPEPEPPKPAPPKQVAPPPRPQVAPHEINAKKS
CKISASRKLHLKILMLGKAKDDLEKEIVDRNEEKEKFLAERVPPLNLSGLSLTDLQNL
CIELHQKIEIVDEERYDFEFKAGKNSYEIHDLSLKILDLRGKFKRPTLRRVRVSADAM
LRALLGSKHKVSMDLRANLKSVKKDDTEKERNVEVSDWRKNVEAKSGMEGRKKM
FDAASAQ

**cu|M00484|GSSSX_GLAND, 26,734.5 Da**
**greenlandsharkSSX (William Joyce (Shiels))**
**45 exclusive unique peptides, 46 exclusive unique spectra, 105 total spectra, 151/233 amino acids (65% coverage)**

MSDQEEQYEE EVIDETEETK EVIEPEPEPP KPAPPKQVAP PPRPQVAPHE INAKKSCKIS
ASRKLHLKIL MLGKAKDDLE KEIVDRNEEK EKFLAERVPP LNLSGLSLTD LQNLCIELHQ
KIEIVDEERY DFEFKAGKNS YEIHDLSLKI LDLRGKFKRP TLRRVRVSAD AMLRALLGSK
HKVSMDLRAN LKSVKKDDTE KERNVEVSDW RKNVEAKSGM EGRKKMFDAA SAQ

>africanlungfishTNNI3
MADEEEVTQYEEEEEYADEEEAEEEEEVKEFEPAKPATPPPPPAAPAPLTRRPSSL
NYRSLVAQPQVKRKSKITASRKLQLKSLMLQIAKQELEREAEERAEEKERYLTQRC
EPLQLSGFSLVELQDLCKQLHATVDVADEERYDLEAKVSKNVQEIEDLNQKIFDLRG
KFKLPQLRRVRMSADAMLRALLGSKHKVCMDLRANLKQVKKDDVEKEIREVGDWR
KNIDAMAGMEGRKKKFEFSLSGQA*

**cu|M00482|ALFC_LFISH, 28,632.0 Da**
**africanlungfishC (William Joyce (Shiels))**
**97 exclusive unique peptides, 121 exclusive unique spectra, 166 total spectra, 205/248 amino acids (83% coverage)**

MADEEEVTQY EEEEEYADEE EAEEEEEVKE FEPAKPATPP PPPAAPAPLT RRPSSLNYRS
LVAQPQVKRK SKITASRKLQ LKSLMLQIAK QELEREAEER AEEKERYLTQ RCEPLQLSGF
SLVELQDLCK QLHATVDVAD EERYDLEAKV SKNVQEIEDL NQKIFDLRGK FKLPQLRRVR
MSADAMLRAL LGSKHKVCMD LRANLKQVKK DDVEKEIREV GDWRKNIDAM AGMEGRKKKF
EFSLSGQA

**Supplementary Figure S1.** Mass spectrometry results showing transcriptomic-predicted *TNNI5* and *TNNI3* sequences from sharks and lungfish, respectively, annotated to show peptide matches (highlighted in yellow) and corresponding protein identification coverage.

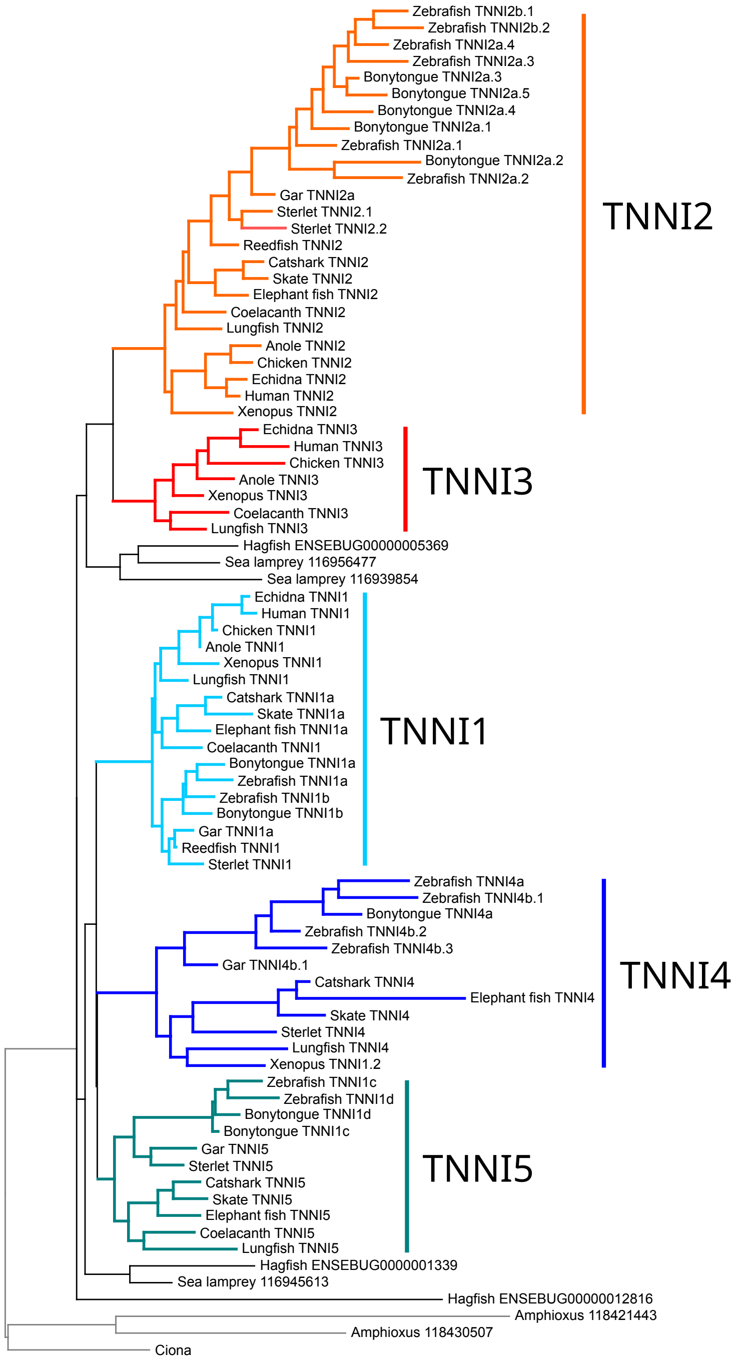


**Supplementary Figure S2.** TNNI phylogenetic tree constrained to show TNNI4-TNNI5 sister relationship. The tree was not significantly different from the maximum likelihood tree (Fig. 1) which showed TNNI1-TNNI5 as sister (Supplementary Table 1).

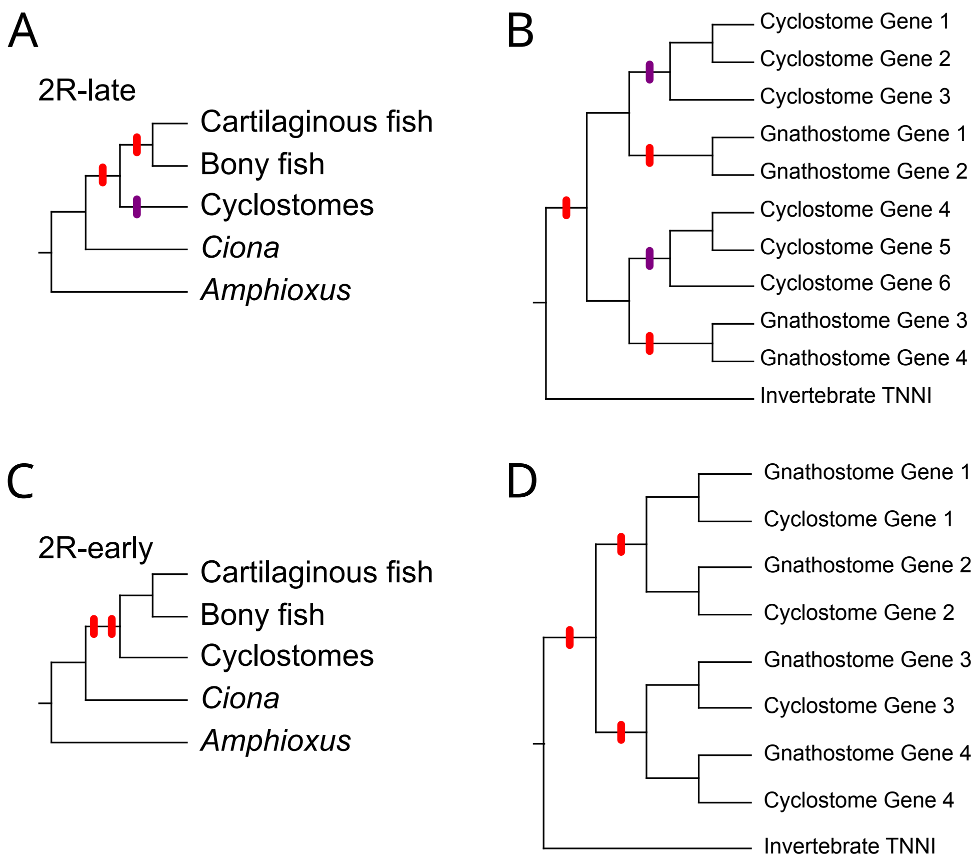


**Supplementary Figure S3.** Alternative hypotheses regarding the timing of the two rounds of whole genome duplication early in vertebrate evolution and their predictions regarding phylogenetic relationships the descending paralogs. There is consensus that gnathostomes underwent two rounds of WGD (Meyer & Schartl 1999; McLysaght et al. 2002; Dehal & Boore 2005), 1R and 2R (shown as red bars), and that 1R predates the split of cyclostomes and gnathostomes. However, the placement of 2R on the vertebrate tree is debated (Kuraku et al. 2009). Recent studies place 2R in the last common ancestor of gnathostomes (Simakov et al. 2020; Nakatani et al. 2021), *i.e.* '2R-late', and suggest that cyclostomes underwent an independent polyploidization early in their evolution (Mehta et al. 2013; Nakatani et al. 2021). By contrast, some authors place 2R in the common ancestor of cyclostomes and gnathostomes (Sacerdot et al. 2018), i.e. '2R-early'. Panels A and C show the placement of these polyploidizations on the vertebrate tree. Panels B and D show the competing phylogenetic predictions.
